# A Universal 6iL/E4 Culture System for Deriving and Maintaining Embryonic Stem Cells Across Mammalian Species

**DOI:** 10.1101/2025.05.20.654948

**Authors:** Duo Wang, Hao Ming, Dongshan Yang, Li-Kuang Tsai, Zhuying Wei, Giovanna Nascimento Scatolin, Xiukun Wang, Kimberly Yau, Litao Tao, Xinyi Tong, Shuling Wang, Kai-Xuan Shi, Denis Evseenko, Ben Van Handel, Bingjing Zhang, Yinjuan Wang, Rajan Iyyappan, Oscar Alejandro Ojeda-Rojas, Guang Hu, Lynda McGinnis, Richard Paulson, Daniel Mckim, Xiangbo Kong, Xiaofeng Xia, Jifeng Zhang, Y. Eugene Chen, Jie Xu, Zongliang Jiang, Qi-Long Ying

## Abstract

The derivation of authentic embryonic stem cells (ESCs) from diverse mammalian species offers valuable opportunities for advancing regenerative medicine, studying developmental biology, and enabling species conservation. Here, we report the development of a robust, serum-free culture system, termed 6iL/E4 that enables the derivation and long-term self-renewal of ESCs from multiple mammalian species, including mouse, rat, bovine, rabbit, and human. Using systematic signaling pathway analysis, we identified key regulators—including GSK3α, STAT3, PDGFR, BRAF, and LATS—critical for ESC maintenance across species. Additionally, inducible expression of KLF2 and NANOG enhances the naive pluripotency and chimeric potential of bovine ESCs. The E4 medium also supports stable ESC growth while minimizing lineage bias. These findings reveal conserved principles underlying ESC self-renewal across divergent mammalian species and provide a universal platform for cross-species stem cell research, disease modeling, and biotechnology applications.

**In Brief:** Wang et al. developed 6iL/E4, a serum-free system sustaining ESCs from mouse, rat, bovine, rabbit, and human. These findings reveal conserved fundamental mechanisms governing ESC self-renewal across diverse mammalian species.

**Highlights:**

- Developed 6iL/E4 system for ESC derivation across five mammalian species.
- PDGFR signaling inhibition as critical for ESC derivation across species.
- E4 medium improves ESC maintenance and avoids neural bias of traditional N2B27.
- Inducible KLF2/NANOG enhances naive pluripotency and chimera formation in bovine.

## INTRODUCTION

Embryonic stem cells (ESCs) are pluripotent stem cells (PSCs) derived from the inner cell mass (ICM) of preimplantation embryos. They have the capacity for long-term self-renewal and can differentiate into all cell lineages both *in vitro* and in chimeric models.^1,2^ However, to date, ESCs capable of contributing to chimeras and transmitting through the germline, which are considered hallmarks of naïve pluripotency, have only been successfully derived from mice^1,2^ and rats^3,4^. During this decades-long effort, the establishment of culture conditions utilizing mitotically inactivated mouse embryonic fibroblasts (MEFs) as feeders and/or leukemia inhibitory factor (LIF), in combination with fetal bovine serum (FBS) or bone morphogenetic protein (BMP), has enabled the successful derivation of ESCs from mice.^5,6^ The development of the well-defined 2i culture condition, consisting of CHIR-99021 (CHIR), a pan-GSK3 inhibitor, and PD0325901(PD03), a MEK1/2 inhibitor, has significantly enhanced the efficiency of deriving and maintaining ESCs from all tested mouse strains.^7^ Unlike earlier culture systems based on LIF plus serum or BMP, which were effective only in a few mouse strains and failed to support rat ESCs (rESCs), the development of 2i or 2i/LIF conditions enabled the successful establishment of rESCs.^3,4^ However, these conditions do not support the derivation of ESCs from other mammalian species. It remains unclear whether a universal culture condition can be developed to support the derivation and maintenance of ESCs across rodents, humans, and other mammalian species.

Among mammalian species, the rabbit serves as a well-established model organism due to its short gestation period (30–31 days), large litter size (4–12 per litter), and suitability for indoor housing.^8^ Phylogenetically, rabbits are closer to humans than mice.^9^ Their anatomical, physiological, genetic, and biochemical similarities to humans make them particularly valuable for studies in pulmonary, cardiovascular, and metabolic research.^9^ Similarly, bovine is not only considered as a highly informative large mammalian model for studying human early embryonic development, given the striking similarities in pre-implantation embryo characteristics,^10–12^ but also hold substantial agricultural and economic value. ESCs from these species represent powerful tools in both basic and translational research as well as genetic engineering and regenerative medicine. However, authentic ESCs from rabbit and bovine (rabESCs and bESCs) analogous to rodent ESCs have yet to be successfully derived. Most reported ESC lines fail to meet the rigorous criteria for pluripotency and therefore cannot be considered true ESCs. For example, rabbit ESCs have been described, but they lack the ability to contribute to chimeric embryos and are often referred to as "ES-like" cells.^13–15^ The established bovine ‘prime’ ESCs^16^ and expanded pluripotent stem cells (EPSCs)^17^ from bovine embryos also have not demonstrated the ability to contribute to chimera formation *in vivo*. These limitations stem in part from suboptimal culture conditions and a lack of understanding of the signaling pathways and molecular mechanisms supporting stem cell maintenance in these species.

We hypothesized that, analogous to the compatibility of culture conditions for both rESCs and mESCs (i.e., 2i or 2i/LIF),^3,4,7^ the core mechanisms governing ESC self-renewal are conserved across mammalian species. While current data suggest species-specific differences, for example, the opposing roles of nuclear β-catenin in rodent versus human ESCs, the underlying signaling networks may share conserved elements that remain to be fully elucidated. In rodents, several key signaling pathways including LIF/STAT3, WNT, and FGF/MAPK have been shown to regulate ESC self-renewal, yet their roles in ESC cultures from other mammals are not well defined. Notably, nuclear β-catenin promotes self-renewal in rodent ESCs but induces differentiation in naive human ESCs (hESC), highlighting the apparent divergence in signaling outcomes.^4,7,18^ However, we believe that these divergent phenotypes may result from differences in pathway context or regulatory feedback, rather than a complete lack of conservation. Supporting this notion, we recently discovered that selective inhibition of GSK3α using BRD0705 promotes the self-renewal of both pluripotent and adult stem cells through a β-catenin-independent mechanism,^19^ suggesting the existence of conserved yet previously unrecognized signaling pathways. Building on these findings, we sought to develop a universal culture condition for ESC derivation and maintenance across multiple species by targeting conserved signaling pathways. We systematically investigated the roles of the LIF/STAT3, WNT, and FGF/MAPK pathways in the derivation of ESCs from rabbit and bovine embryos. As a result, we developed a serum-free ‘6iL/E4’ culture system that enabled the successful generation of rabESCs and bESCs. We further demonstrated that the ‘6iL/E4’ system supports chimera- and germline-competent mESCs and naive rESCs, as well as naive hiPSCs and hESCs. Together, our findings provide compelling evidence for a conserved signaling framework governing ESC self-renewal and establish a universal culture system applicable to multiple mammalian species.

## RESULTS

### Development of the ‘6iL/E4’ Culture Medium Enables the Derivation of Rabbit and Bovine ESCs

An effective ESC culture system typically includes a basal medium combined with selected small molecules and/or growth factors to support self-renewal. To date, only the 2i/N2B27 system has been validated as a universal method for deriving both mESCs and rESCs.^3,4,7^ We initially applied 2i/N2B27 to derive ESCs from bovine and rabbit embryos; however, by day 5, all bovine and rabbit embryo-derived cells underwent complete differentiation (Figure S1A). This suggested that certain components within the 2i/N2B27 system may trigger differentiation in non-rodent species. Given the complex composition of N2B27 and its known ability to drive neural differentiation of mESCs in the absence of 2i,^20^ we hypothesized that some N2B27 additives may be detrimental to ESC maintenance. To address this, we systematically assessed the necessity of each N2B27 component in supporting mESC self-renewal under 2i conditions. Using the DMEM-F12/Neurobasal basal medium supplemented with only insulin, transferrin, and BSA, the three key components of N2B27, we tested the effects of reintroducing each omitted additive. This screening revealed that sodium selenite was the only additional component required for long-term expansion of mESC (Figures S1B and S1C). We termed this minimal formulation ‘E4’ medium and confirmed that 2i/E4 robustly supported long-term self-renewal of both mESCs and rESCs (Figures S1D and S1E). Notably, rESCs cultured in 2i/E4 exhibited enhanced colony formation and reduced cell death (Figure S1E). Unlike N2B27, The E4 medium did not induce neural differentiation of ESCs in the absence of 2i, highlighting its specificity in supporting ESC self-renewal (Figure S1F). Thus, E4 serves as a simplified and optimized basal medium that excludes extraneous additives potentially responsible for promoting unintended lineage commitment.

We next screened small molecules and growth factors to identify conditions enabling ESC derivation from bovine and rabbit embryos. Recognizing the reliance of rodent ESCs on WNT activation and FGF pathway inhibition for naïve pluripotency,^3,4,7^ we began by probing the role of WNT signaling in the derivation of ESCs from bovine and rabbit embryos. In rodent ESCs, GSK3 inhibition stabilizes β-catenin to promote nuclear translocation and self-renewa.^7^ However, recent studies in naive human ESCs showed that activating β-catenin instead promotes differentiation and cell death.^18^ Consistent with this, CHIR induced differentiation in both bovine and rabbit inner cell mass (ICM) outgrowths (Figure 1A). In contrast, BRD0705, a GSK3α-specific inhibitor, combined with the WNT pathway inhibitor IWR1 suppressed non-ESC-like cell expansion more effectively than CHIR (Figure 1A), suggesting its suitability for maintaining pluripotency. Nevertheless, BRD0705/IWR1 alone could not sustain passaged ESCs (data not shown), indicating the need for additional factors.

**Figure 1.**
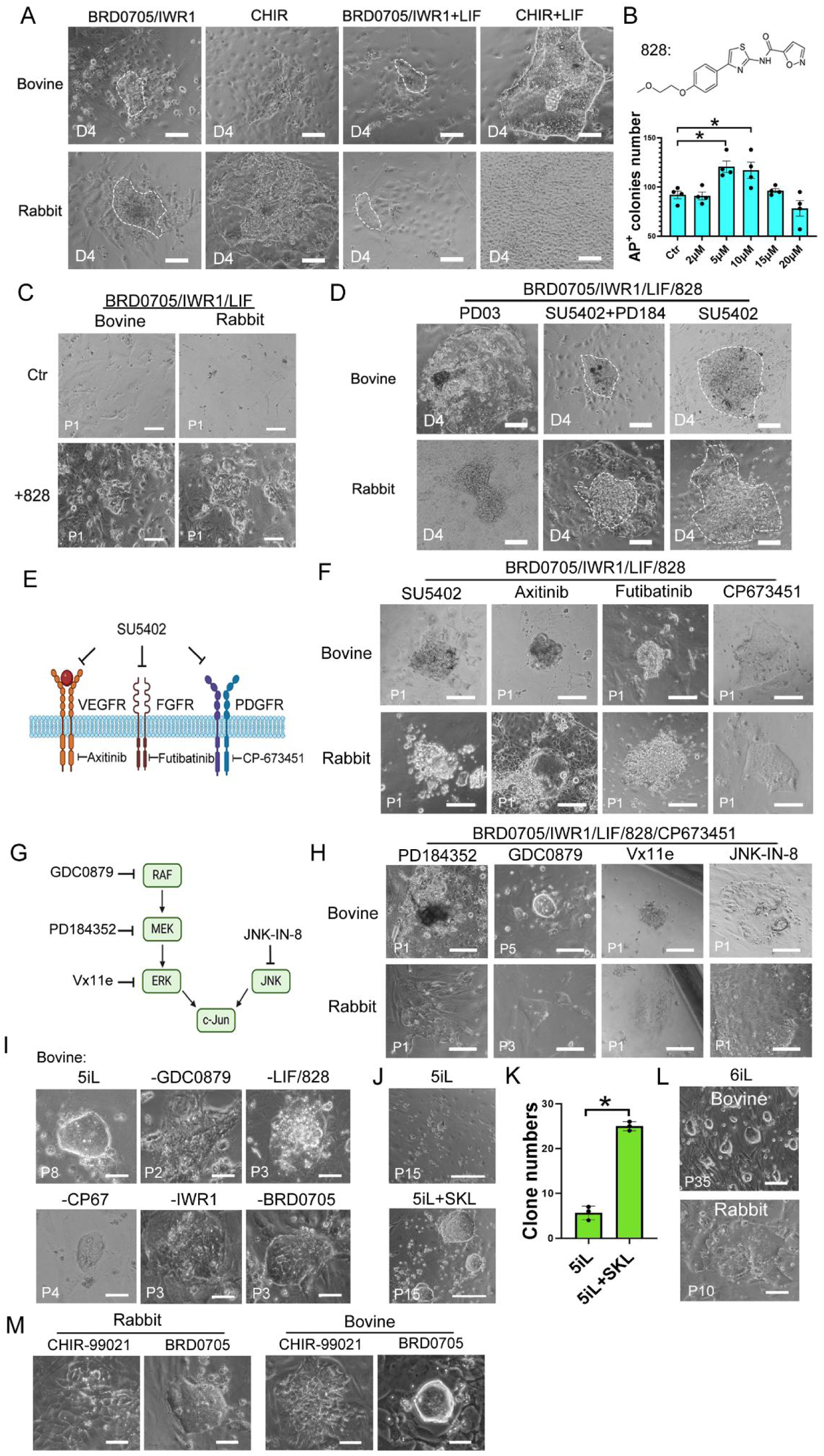
Optimization and Evaluation of Culture Conditions for Deriving and Expanding ESCs from bovine and rabbit. (A) Representative phase-contrast images of bovine and rabbit ICMs cultured under different conditions on day 4. Scale bars, 100 μm. Dashed outlines indicate undifferentiated ICM outgrowths. (B) Chemical structure of compound 828 and quantification of AP+ colonies in mESC cultures treated with 20 ng/ml LIF and varying concentrations of 828. Data are presented as mean ± SEM, with statistical significance indicated (*P < 0.05). (C) Representative Passage 1 (P1) cell morphologies from bovine and rabbit ICMs cultured under BRD0705/IWR1/LIF with or without 828. Scale bars, 100 μm. (D) Representative outgrowth morphology from bovine and rabbit ICMs cultured for 4 days under BRD0705/IWR1/LIF/828 conditions with specific signaling pathway inhibitors. Scale bars, 100 μm. Dashed outlines indicate undifferentiated ICM outgrowths. (E) Schematic illustration of the targets of SU5402. (F) Representative cell morphology of P1 ESCs derived from bovine and rabbit embryos and cultured with SU5402, Axitinib, Futibatinib, or CP-673451 in BRD0705/IWR1/LIF/828 E4 medium. Scale bar, 100 µm. (G) Diagram of the MAPK signaling cascade with small-molecule inhibitors used in this study: GDC0879 (RAF), PD184352 (MEK), Vx11e (ERK), and JNK-IN-8 (JNK). (H) Phase-contrast images showing the effects of MAPK pathway inhibitors on bESCs and rabESC derivation. ESCs were cultured in E4 medium supplemented with BRD0705, IWR1, LIF, 828, and CP67, with individual MAPK pathway inhibitors added separately. Scale bar, 100 µm. (I) Representative images of bESCs derived from bovine blastocysts and cultured in 5iL E4 medium, with or without the omission of individual components. Scale bars, 50 µm. (J) Phase-contrast images of bovine ESCs cultured under 6iL conditions, consisting of 5iL medium supplemented with the additional small molecule SKL. Scale bars, 200 µm. (K) Quantification of bESC colony numbers under ‘5iL’ medium with or without small molecules compound SKL. Data are presented as mean ± SEM. * p < 0.05. (L) Phase-contrast images of P35 bovine ESCs derived from blastocysts and P10 rabESCs derived from morula stage embryos in ‘6iL’ E4 medium. Scale bars = 100 µm. (M) Phase-contrast images of bESCs and rabESCs derived in 5iL plus CHIR or BRD0705. Scale bars = 50 µm.

We also examined the STAT3 pathway. Adding LIF to the BRD0705/IWR1 combination promoted the formation of more ESC-like colonies from ICM outgrowths in both rabbit and bovine embryos (Figure 1A). To improve efficacy and universality, we screened additional STAT3 activators and identified a novel small molecule, 828,^21^ which enhanced mESC colony formation when combined with LIF (Figures 1B, S2A and S2B). However, the combination of BRD0705/IWR1/LIF/828 produced colonies with vacuole formation in bovine and rabbit cultures, leading to eventual differentiation and death (Figure 1C).

To further refine the system, we examined FGF/MAPK pathway inhibition based on the BRD0705/IWR1/LIF/828 regimen. We tested PD03 (used in 2i),^7^ SU5402, and PD184352 (PD184) (used in the ‘3i’ system).^7^ SU5402 effectively suppressed non-ICM outgrowths and enhanced ESC-like outgrowths from both bovine and rabbit ICMs (Figure 1D). As SU5402 targets VEGFR, FGFR, and PDGFR pathways, we subsequently tested specific inhibitors: Axitinib (VEGFR), Futibatinib (FGFR), and CP673451 (PDGFR) (Figure 1E). Only CP673451 (CP67) promoted stable ESC colony formation after passaging (Figure 1F). CP67 and SU5402 also enhanced rESC colony numbers when combined with CHIR and PD184, implicating PDGFR inhibition as a key contributor (Figures S2C and S2D). Based on these results, we refined the culture to BRD0705/IWR1/LIF/828/CP67. However, this supported only limited passaging (2 passages) of rabbit and bovine ESCs. We further screened MAPK inhibitors, including GDC0879 (a B-Raf inhibitor), PD184 (a MEK inhibitor), Vx11e (an ERK inhibitor), and JNK-IN-8 (a JNK inhibitor) (Figure 1G). We found GDC0879 significantly enhanced passaging capacity of both rabbit and bovine ESC-like colonies (Figure 1H). This optimized six-component regimen (BRD0705, IWR1, LIF, 828, CP67, and GDC0879) was designated ‘5iL’. Subtractive experiments confirmed that removal of any single component abolished bESC self-renewal (Figure 1I).

Although the ‘5iL’ system enabled efficient derivation of ESCs from both rabbit and bovine embryos, bESCs exhibited reduced proliferation after ∼15 passages. To address this limitation, additional small molecules were screened, and this identified the Wnt/β-catenin pathway inhibitor SKL2001(SKL)^22^ that significantly increased the efficiency of bESC colony formation and enabled rabESC maintenance (Figures 1J and 1K). This led to the final optimized ‘6iL’ formulation: BRD0705, IWR1, CP67, GDC0879, SKL2001, 828, and LIF in E4 medium for the derivation of bovine and rabbit ESCs (Figure 1L).

Lastly, replacing BRD0705 with CHIR in this optimized system caused widespread differentiation in both species (Figure 1M), reinforcing that ESCs from non-rodent mammals respond differently to WNT activation and that selective GSK3α inhibition is essential. Taken together, these results establish ‘6iL/E4’ as a robust culture system for deriving and maintaining ESCs from rabbit and bovine embryos.

### Stable and Long-Term Maintenance of Naïve Rodent ESCs Cultured in 6iL

To validate the universality of the 6iL culture condition, we tested its ability to support naive ESC derivation and maintenance in rodent species. Using mouse blastocysts, we successfully derived mESC lines under 6iL conditions that could be stably passaged for over 25 generations (Figure 2A). These 6iL-mESCs exhibited normal karyotypes (Figure S3A), retained alkaline phosphatase (AP) activity (Figure 2A), and expressed core pluripotency markers NANOG and OCT4 (Figure 2B). Furthermore, they maintained the capacity to differentiate into derivatives of all three germ layers in vitro (Figure 2C). Although RNA-seq results revealed that 6iL-mESCs exhibited a unique gene expression profile compared to 2iL-mESCs and AFX-EpiSCs (Figure S3B), qRT-PCR analysis of PSC-related gene expression showed that 6iL-mESCs are distinct from EpiSCs and FS cells. The expression levels of *Tfcp2l1*, *Rex1*, and *Sox2* in 6iL-mESCs were comparable to those of naive mESCs cultured in 2iL (Figure S3C). Unlike EpiSCs, 6iL-ESCs did not express *Foxa2* and exhibited lower expression of *T* and *Gata4* (Figure S3C). However, Otx2 expression in 6iL-mESCs more closely resembled that in EpiSCs and FS cells than in naive mESCs. (Figure S3C). To further demonstrate the pluripotency of 6iL-mESCs, we injected GFP-labeled 6iL-mESCs into WT E3.5 mouse blastocysts. GFP+ cells were observed in 7 of 21 embryos at E9.5 (Figures 2D, S3D, and S3E). To validate the chimeric nature of these embryos, we dissociated them into single cells, cultured the resulting cells, and confirmed GFP expression in the viable cell population (Figure 2E). Furthermore, GFP+ embryonic germ cells (EGCs) were derived from the gonads of these embryos under CHIR/PD03/LIF (2iL) conditions^23,24^ and expanded in long-term culture (Figures 2F and 2G). The EGCs expressed OCT4, confirming their identity (Figure 2H). In conclusion, the 6iL condition provides a robust system for deriving and maintaining chimera- and germline-competent mouse ESCs.

**Figure 2.**
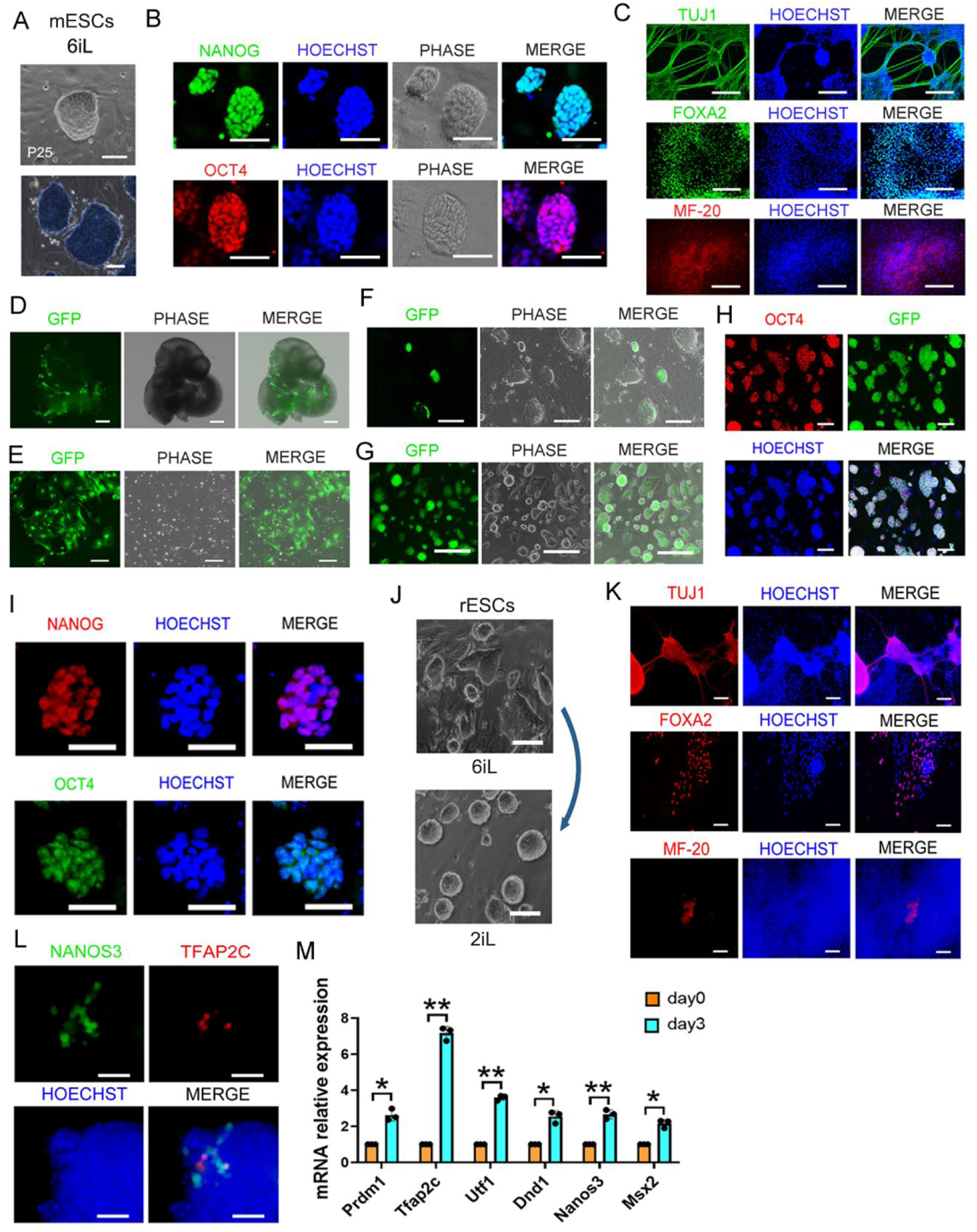
Characterization of mESCs and rESCs derived and maintained in 6iL. (A) Top: Representative phase-contrast images showing ESCs cultured under 6iL conditions. Scale bars = 100 µm. Bottom: AP staining of ESC colonies derived under 6iL conditions. Scale bars = 50 µm. (B) Representative IF images of 6iL-derived ESCs (passage 15). Green indicates NANOG; red indicates OCT4; blue indicates HOECHST. Scale bars = 100 µm. (C) Representative IF images of EB outgrowths for multi-lineage differentiation markers. Scale bars = 200 µm. (D) Representative FL images of E9.5 days chimaeras from blastocyst injected with GFP labeled 6iL-mESCs. Scale bars = 500 µm. (E) Representative FL images of GFP+ cells cultured from dissociated E9.5 chimeric embryos. Scale bars = 100 µm. (F) Representative phase-contrast and FL images of GFP+ clones derived from the gonads of E13.5 chimeric embryos (chimaeras from blastocyst injected with GFP labeled 6iL-mESC) cultured in 2iL. Scale bars = 200 µm. (G) Representative phase-contrast and FL images of passage 3 EGCs derived from the GFP+ mono-clone shown in Figure (F), cultured under 2i/LIF conditions. Scale bars, 200 μm. (H) IF results confirming OCT4(red) expression in the GFP+ EGCs in Figure (G). Blue is Hoechst. Scale bars = 50 µm. (I) Representative IF analysis of rESCs cultured in 6iL, showing expression of pluripotency markers NANOG (red) and OCT4 (green). HOECHST (blue) marks nuclei. Scale bars = 50 µm. (J) Representative images of rESCs cultured in 6iL for six passages, and of 6iL-rESCs following transition to 2i conditions for two additional passages. Scale bars = 100 µm. (K) IF analysis of EB outgrowths derived from 6iL-cultured rESCs, showing expression of lineage-specific markers: TUJ1 (ectoderm), FOXA2 (endoderm), and MF-20 (mesoderm). Nuclei are counterstained with HOECHST (blue). Scale bars = 100 µm. (L) IF analysis of PGC-LCs differentiated from 6iL-rESCs, showing expression of PGC-specific markers NANOS3 (green) and TFAP2C (red). Nuclei are counterstained with HOECHST (blue). Scale bars = 20 µm. (M) qRT-PCR analysis of PGC marker gene expression of PGC-LCs derived from 6iL-cultured rESCs. Data are presented as mean ± SEM. *p< 0.05, **p < 0.01.

Next, we evaluated the ability of 6iL in sustaining rESCs. rESCs initially derived under 2i were passaged in 6iL for at least six passages. Immunofluorescence (IF) analysis demonstrated strong and consistent expression of the pluripotency markers NANOG and OCT4, confirming the maintenance of their undifferentiated state (Figure 2I). Notably, when 6iL-cultured rESCs were transitioned back to the 2i, they remained viable and readily adapted to the naive rESC culture medium (Figure 2J). These findings demonstrated that the 6iL effectively preserved naive rESC identity. Furthermore, *in vitro* embryoid body (EB) differentiation assays confirmed that rESCs maintained in 6iL successfully gave rise to derivatives of all three germ layers (Figure 2K), providing strong evidence of their pluripotency. We also evaluated the capacity of rESCs cultured in 6iL to differentiate into primordial germ cell-like cells (PGC-LCs). IF analysis revealed that these cells could be induced to differentiate into NANOS3 and TFAP2C double-positive cells *in vitro* (Figure 2L), and qRT-PCR analysis confirmed the elevated expression of key PGC marker genes (Figure 2M). Together, these findings demonstrate that 6iL effectively supports the cultivation and maintenance of naive rESCs while preserving their ability to self-renew and differentiate into PGC-LCs and other cell lineages.

### Derivation and Characterization of 6iL-bESCs

Building on the demonstrated reliability of the 6iL condition for culturing mESCs and rESCs, we next characterized the long-term self-renewal capacity of bESCs established under 6iL. Our results demonstrated that the 6iL condition supported the derivation of bESCs from bovine blastocysts (Figure 3A). After long-term expansion under 6iL, bESCs maintained the correct karyotype (Figure S4A). Further characterization revealed that bESCs maintained stable colony morphology after long-term *in vitro* culture (over 35 passages) (Figure 3A) and that bESCs exhibited robust expression of key pluripotency markers, including NANOG and OCT4 (Figure 3B). Others have derived embryonic disc stem cell lines using a media containing Activin A, FGF, and XAV939 (AFX).^25^ Compared with cells maintained in AFX-, 6iL-bESCs exhibited increased expression of naive pluripotency markers, including *Nanog*, *Pou5f1*, and *Rex1*, along with notably lower expression of the formative-stage transcription factor *Otx2*^26,27^ (Figure 3C). *In vitro* differentiation assay demonstrated that 6iL-bESCs were capable of generating derivatives of all three germ layers, evidenced by the expression of markers for endoderm (GATA4), mesoderm (MF-20), and ectoderm (TUJ1) (Figure 3D). In conclusion, 6iL-bESCs exhibit hallmarks of pluripotency.

**Figure 3.**
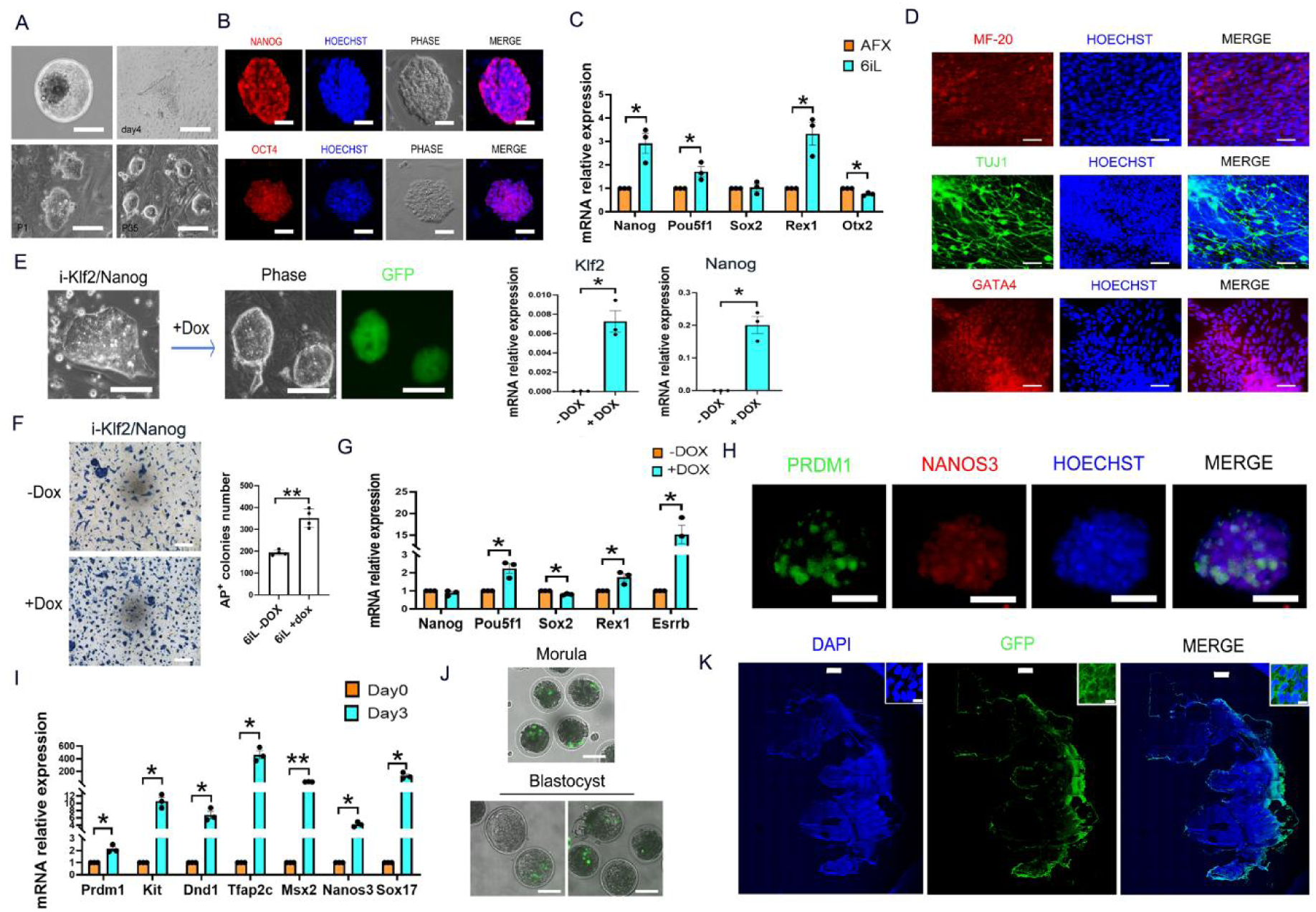
Derivation and Characterization of 6iL-bESCs. (A) Representative images of bovine blastocyst, outgrowths and ESC colonies (passage 1 and 35) derived under the ‘6iL’ condition. Scale bars = 100 µm. (B) IF staining of 6iL-bESCs showing NANOG and OCT4 expression. Scale bars = 50 µm. (C) qRT-PCR comparing pluripotency markers (Nanog, Pou5f1, Sox2, Rex1) and formative marker Otx2 in 6iL-bESCs versus AFX-cultured cells. (mean ± SEM; p< 0.05, ns, not significant). (D) Representative IF images of EB outgrowths demonstrating mesoderm (MF-20), ectoderm (TUJ1), and endoderm (GATA4) differentiation. Scale = 50 µm. (E) Representative Images of GFP-labeled 6iL-bESCs with a DOX-inducible Klf2/Nanog system. Scale bars = 100 µm. The right panel shows the qRT-PCR results demonstrating the expression of exogenous Klf2 and Nanog genes after the addition of DOX. (F) AP staining of i-Klf2/Nanog-expressing bovine ESCs ± DOX under 6iL conditions with corresponding quantification. Scale = 200 µm. (G) qRT-PCR analysis of pluripotency gene expression in 6iL-bESCs ± DOX. (H) IF images of PGC-LC cells derived from 6iL-bESCs showing PRDM1 (green) and NANOS3 (red). Scale bars= 50 µm. (I) qRT-PCR of PGC marker genes during differentiation of 6iL-bESCs into PGC-LCs (mean ± SEM; *p* < 0.05, \**p* < 0.01). (J) Representative images of morula- and blastocyst-stage bovine embryos injected with DOX-inducible GFP-labeled 6iL-bESCs. Scale bars= 100 µm. (K) IF analysis of bovine embryo sections from day 40 chimeras, which were developed from blastocyst embryos injected with DOX-inducible Klf2/Nanog-expressing GFP-labeled bESCs. Scale bars, 100 μm. Insets show enlarged images. Scale bars, 500 μm.

It has been demonstrated that NANOG sustains *Oct4* expression and mESC pluripotency independent of STAT3 signaling.^28^ Klf2 is negatively regulated by MEK/ERK signaling, and its overexpression can maintain mESCs in the ground state.^29^ Transient expression of these two pivotal factors is sufficient to activate the pluripotency network, thereby resetting the human pluripotent state.^30^ Therefore, we introduced doxycycline (DOX)-inducible Klf2/Nanog into GFP-labeled 6iL-bESCs to further sustain its naive pluripotency. Upon DOX induction, bESCs exhibited increased colony compaction (Figure 3E) as well as enhanced proliferation and growth (Figures 3F), accompanied by an upregulated expression of pluripotency genes including *Pou5f1*, *Rex1*, and *Esrrb* (Figure 3G). We next investigated whether 6iL-bESCs possess the potential to generate PGC-LCs. Following DOX withdrawal, the 6iL cells were first treated with Activin A/bFGF overnight, then induced to differentiate into PGC-LCs following the previously reported methods.^17,31^ Differentiation into PGC-LCs was demonstrated by the presence of PRDM1/NANOS3 double-positive cells (Figure 3H) and the upregulation of PGC marker genes (Figure 3I). These findings demonstrated that exogenous induction of Klf2/Nanog did not impair the normal differentiation of ESCs following DOX withdrawal, and that these cells retain the capacity to generate PGC-LCs *in vitro*. Finally, encouraged by these findings, we examined the developmental capacity of bESCs to contribute to the formation of chimeras. We microinjected DOX-inducible Klf2/Nanog-expressing GFP-labeled bESCs into morula-stage or early blastocyst-stage bovine embryos (Figure 3J). Following DOX withdrawal, the embryos were cultured *in vitro* for 24 to 48 hours. GFP-positive cells were detected in 102 of 150 morulae and 95 of 150 blastocysts (Figures 3J, S4B-D). Of these, blastocysts with clear GFP signals were transferred to 10 recipient cows, that yielded 8 surrogates with confirmed pregnancy. Immunostaining analysis of day 40 bovine embryos revealed that GFP-positive cells were detected in one chimeric embryo (Figure 3K). These results demonstrated that bESCs derived under 6iL conditions, combined with inducible Klf2/Nanog expression, were capable of generating bESC-derived bovine chimeric embryos *in utero*. Thus, the 6iL culture system may serve as a powerful tool for studying embryo formation in large mammals and for bovine genetic engineering. Furthermore, this is conceptually important because it demonstrates fundamentally conserved mechanisms governing ‘ground state’ stem cell pluripotency across mammals with divergent phylogeny and body size.

### Derivation and Characterization of rabESCs in 6iL/TDI

Next, we aimed to apply the 6iL condition to derive rabbit ESCs. We observed that a subset of rabESC clones underwent differentiation after 10 passages under the 6iL condition (Figure 4A). To address this, we screened additional small molecules and identified the LATS1/2 inhibitor TRULI and its improved derivative, TDI-011536 (TDI),^32^ as promising candidates to improve the maintenance of rabESCs (Figure S5A). The addition of either TDI or TRULI to the ‘6iL’ greatly enhanced the long-term expansion of rabbit ESCs (Figure 4A). Thus, 6iL+TDI provided the most optimized culture condition for rabESCs. Interestingly, replacing the WNT/β-catenin inhibitor SKL with the LATS1/2 inhibitors (TRULI and TDI) in bESC cultures resulted in similar or improved bESC expansion (Figures 4B and 4C). Similar effects were also observed in mouse and rat ESCs (data not shown). Therefore, TDI or TRULI exhibits a universal effect on the culture of ESCs from divergent species and can serve as an alternative to WNT/β-catenin inhibition by SKL.

**Figure 4.**
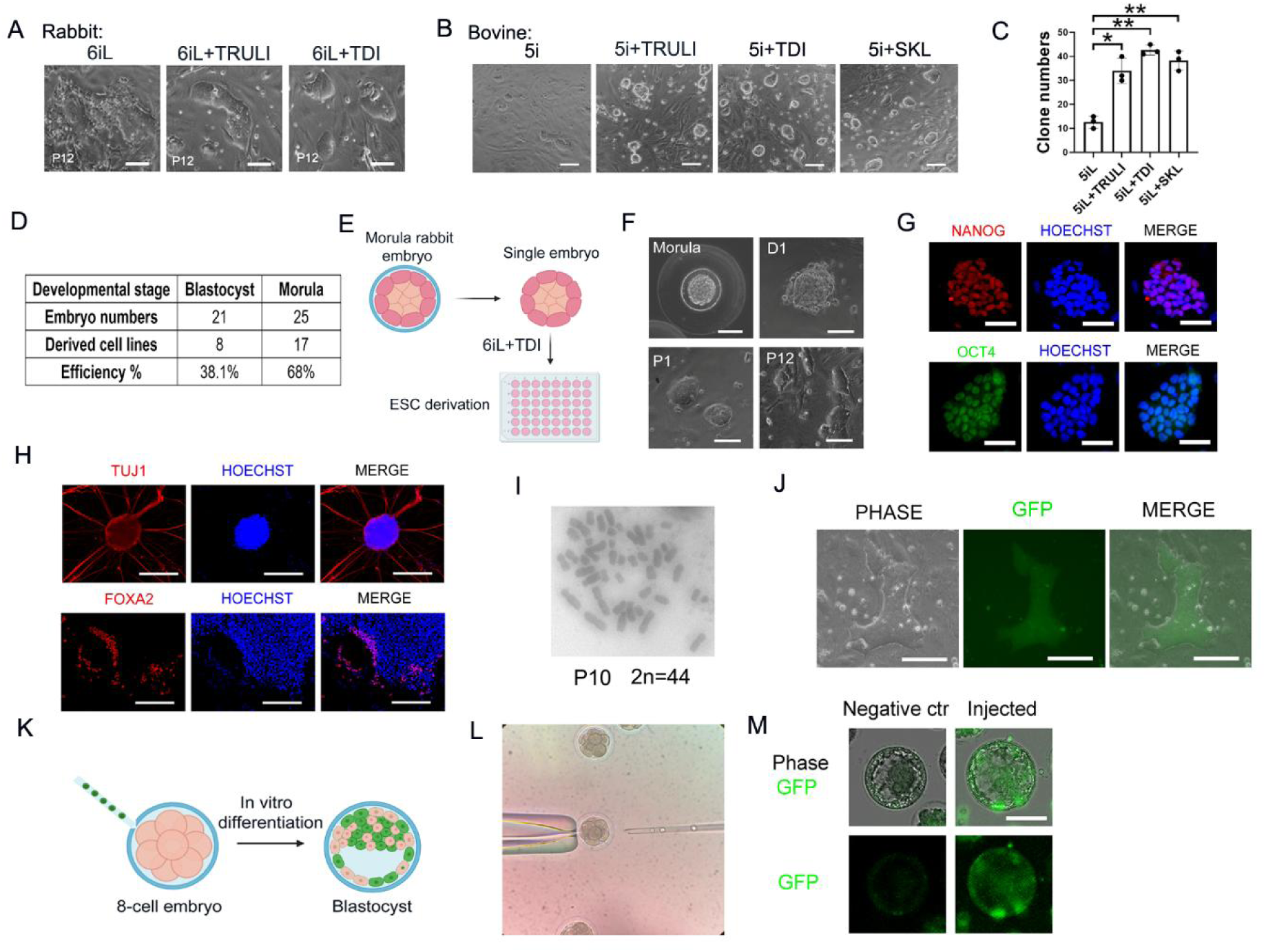
Derivation and Characterization of rabESCs in 6iL/TDI. (A) Representative phase-contrast images of rabESCs derived from morula-stage embryos in 6iL, 6iL+TRULI, or 6iL+TDI in E4 medium at P12. Scale bars = 100 µm. (B) Representative phase-contrast images of bESCs cultured under ‘5iL’ conditions with additional small molecules (TRULI, TDI, and SKL). Scale bars = 200 µm (C) Quantification of bESC colony numbers under ‘6iL’ conditions with additional small molecules. Data are presented as mean ± SEM. * p < 0.05, ** p < 0.01. (D) Summary table showing the efficiency of rabESC derivation from different developmental stages. (E) Schematic representation of rabESC derivation from individual morula embryos under 6iL conditions. (F) Representative phase-contrast images showing the morphology of rabESCs derived from the morula stage. Scale bars = 100 µm. (G) Representative IF images of 6iL-TDI-derived rabESCs showing robust expression of pluripotency markers NANOG (red) and OCT4 (green). Scale bars = 50 µm. (H) Representative IF images of EBs differentiation derived from 6iL rabESCs, confirming expression of lineage-specific markers: TUJ1 (ectoderm, red), FOXA2 (endoderm, red), and HOECHST (blue). Scale bars = 100 µm. (I) Karyotype analysis of passage 10 (P10) rabESCs, showing a normal diploid chromosome number (2n=44). (J) Representative phase-contrast and FL images showing GFP-labeled rabESC colonies derived under 6iL conditions. Scale bars = 100 µm. (K) Schematic of generating chimeric rabbit embryos by transferring GFP-labeled rabESCs into 8-cell embryos that develop into blastocysts *in vitro*. (L) Microinjection of GFP-labeled 6iL ESCs into 8-cell stage rabbit embryos. (M) Representative FL images showing the contribution of GFP+ rabESCs to rabbit chimeric embryos. Ten GFP-labeled rabESCs were injected into each 8-cell stage rabbit embryos, and fluorescence images show successful integration into blastocysts. Scale bars: 100 μm.

Subsequently, we derived and cultured rabESCs under 6iL+TDI. We found that 6iL + TDI enabled more efficient derivation of rabESCs from morula-stage embryos compared to blastocyst-stage embryos (Figure 4D). In combination with 6iL, the addition of SKL and TDI (or TRULI) significantly enhanced rabESC colony formation and facilitated their stable, long-term passaging (Figures 4E and 4F). The pluripotency of 6iL-rabESCs was validated by IF analysis, which demonstrated robust expression of the pluripotency markers NANOG and OCT4 (Figure 4G). *In vitro* EB differentiation assays demonstrated that 6iL-rabESCs differentiated into derivatives of all three germ layers—ectoderm (Figure 4H), mesoderm (Movie S1), and endoderm (Figure 4H). Karyotype analysis confirmed that 6iL-rabESCs maintained a normal chromosome complement (2n = 44) after long-term culture (Figure 4I). Additionally, we generated GFP-labeled 6iL-rabESCs (Figure 4J) and assessed their integration potential by microinjecting 10 GFP-labeled 6iL-rabESCs into each 8-cell stage host embryos from non-GFP animals (Figures 4K and 4L). We observed that the injected embryos developed into blastocysts with a high degree of chimerism, with GFP-positive cells detected in both the ICM and trophectoderm (Figures 4M and S5B). These data demonstrate that the 6iL/TDI culture system enables efficient derivation of rabESC lines from morula-stage embryos.

### Generation and Characterization of Human PSCs using 6iL

Building upon the successful derivation and expansion of ESCs from bovine, rabbit, mouse, and rat using 6iL, we investigated its applicability to human PSCs. Murine induced PSCs (iPSCs) derived from somatic cells via overexpression of Oct4, Sox2, Klf4, and c-Myc have achieved full pluripotency under mESCs culture conditions, enabling the generation of full-term mice through tetraploid complementation.^33^ These findings underscore that overexpression of key transcription factors can reprogram somatic cells into a naive state, provided that appropriate culture conditions preserve pluripotency for chimera formation. We generated integration-free human iPSCs (hiPSCs) by reprogramming human umbilical cord blood cells using plasmids encoding SOX2, KLF4, OCT3/4, L-MYC, and LIN28a.^34^ Following electroporation, cells were transferred to feeder-coated plates on day 3, and the medium was replaced with 6iL (Figure 5A). By day 12, multiple iPSC-like colonies emerged (Figure 5B). Individual colonies were isolated and passaged with stable long-term self-renewal capacity under 6iL (Figure 5C). Notably, 6iL-hiPSCs tolerated single-cell passaging with 0.025% trypsin, obviating the need for ROCK inhibitors. Further characterization revealed that 6iL-hiPSCs exhibited robust expression of pluripotency markers NANOG and OCT4 (Figure 5D) and maintained a normal karyotype (2n = 46) (Figure S6A). *In vitro* EB differentiation assays demonstrated the three germ layers differentiation potential of 6iL-bESCs (Figure 5E). Principal component analysis (PCA) of the transcriptomes of 6iL -hiPSCs with 4iWIS2-hESCs,^35^ HNES3-hESCs,^36^ and reset H9-hESCs^30^ positioned 6iL-hiPSCs between reset H9-hESCs and 4iWIS2-hESCs, further supporting their pluripotency (Figure 5F). Interestingly, we found that 6iL-hiPSC can be directly induced to differentiate into PGC-LCs. IF analysis demonstrated the expression of PGC-specific markers, including TFAP2C, PRDM1, and NANOS3, in the induced PGC-LCs (Figures 5G and S6B). qRT-PCR analysis further corroborated these findings by showing a significant upregulation of PGC marker genes relative to baseline levels at the onset of differentiation (Figure 5H).

**Figure 5.**
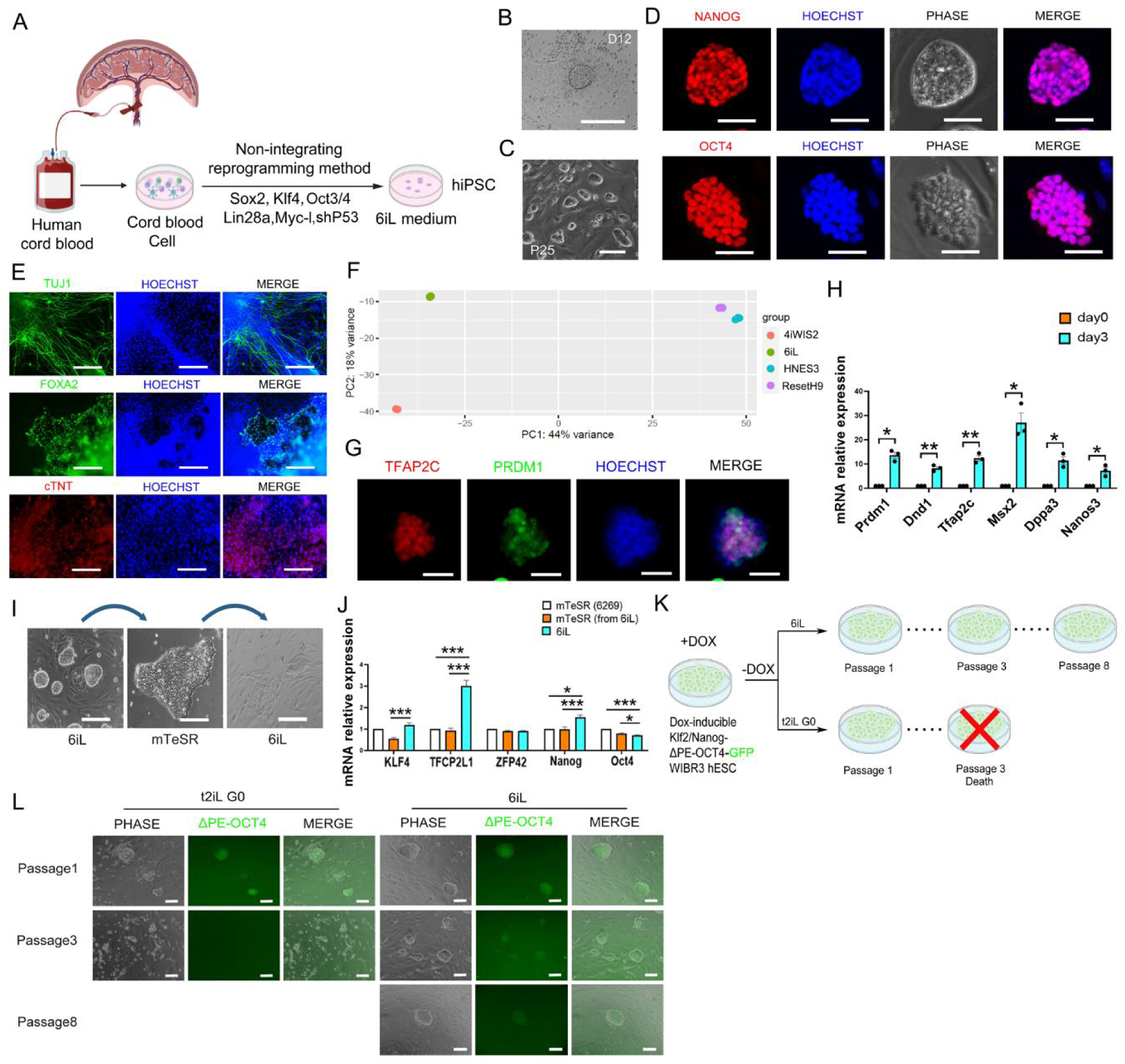
Generation and Characterization of Human PSCs using 6iL. (A) Schematic of the reprogramming strategy for generating hiPSCs. (B) Representative phase-contrast image of a hiPSC clone derived from human cord blood cells at day 12 of reprogramming. Scale bar = 200 µm. (C) Representative images of hiPSC cultured under the 6iL condition at passage 25. Scale bar = 100 µm. (D) IF analysis confirming expression of the naive pluripotency markers NANOG and OCT4 in 6iL-hiPSCs. HOECHST (blue) marks nuclei. Scale bars = 50 µm. (E) Representative IF images of EB outgrowths derived from 6iL-hiPSCs, showing expression of lineage-specific markers: TUJ1 (ectoderm, green), FOXA2 (endoderm, red), and cTNT (mesoderm, red). Scale bars = 200 µm. (F) A PCA plot of RNA-seq data from 6iL-hiPSC and hESC cell lines established under different representative hESC culture conditions. (G) IF analysis of PGC-LCs differentiated from 6iL-hiPSCs, showing expression of PGC-specific proteins TFAP2C (red), PRDM1 (green), and HOECHST (blue) marks nuclei. Scale bars = 50 µm. (H) qRT-PCR analysis of PGC marker gene expression on days 0 and 3 of PGC-LCs differentiated from 6iL-hiPSCs. Data are presented as mean ± SEM. *p < 0.05, **p < 0.01. (I) Cell morphology of hiPSCs cultured under 6iL conditions; their morphology after transitioning to mTeSR conditions; and the morphology of cells cultured for 7 passages in mTeSR conditions and subsequently reverted to 6iL+Go/feeder conditions. Scale bars = 200 µm. (J) qRT-PCR analysis comparing the expression of pluripotency marker genes in hiPSCs cultured under 6iL conditions, cells transitioned to mTeSR conditions, and the 6269 hiPSC cell line cultured in mTeSR conditions. Data are presented as mean ± SEM. *p < 0.05, **p <0.01, ***p<0.001. (K) Schematic of the ΔPE-Oct4-GFP reset system in WIBR3 hESCs. Cells were transfected with ΔPE-Oct4-GFP and subjected to a DOX-inducible Klf2/Nanog system to induce a naive state. After DOX withdrawal, cells were cultured under 6iL or t2iL Go conditions. (L) Representative FL images of ΔPE-Oct4-GFP+ WIBR3 hESCs with a DOX-inducible Klf2/Nanog system. After DOX withdrawal, the cell morphology was observed under 6iL and t2iL Go conditions at passages 1, 3, and 8. Scale bars = 100 µm.

The differences between 6iL-hiPSCs and hiPSCs cultured in mTeSR, a widely used commercial medium, were also characterized. We found that transitioning 6iL-hiPSCs from 6iL into mTeSR resulted in a morphological shift from domed to flattened colonies (Figure 5I). However, reverting hiPSCs cultured in mTeSR for 7 passages back to 6iL led to significant cell death and differentiation (Figure 5I), demonstrating that the states of these two cell types are divergent. Further qRT-PCR analysis showed that 6iL-hiPSCs exhibited significant higher expression of pluripotency markers *Tfcp2l1* and *Nanog* compared to mTeSR-cultured hiPSCs (Figure 5J). Similar results were obtained with the commercial hiPSC line 6269, indicating that hiPSCs cultured under 6iL conditions represent an earlier developmental stage than those maintained in mTeSR medium. Thus, 6iL supports maintenance of widely used hiPSC lines in a more primitive and naïve pluripotent state.

To further evaluate the capacity of 6iL to sustain naive hESCs, WIBR3 hESCs were transfected with a ΔPE-Oct4-GFP and introduced into a DOX-inducible Klf2/Nanog system following the established protocol.^30^ Induction of *Klf2* and *Nanog* via DOX reset the hESCs to a more naive state, as confirmed by ΔPE-Oct4-GFP expression (Figure S6C). Following DOX withdrawal, the cells were cultured and passaged under either 6iL or the previously reported t2iL/Go-naive ESC culture condition^30,36^ (Figure 5K). Under t2iL/Go conditions, most cells died by the third passage, whereas cells cultured in 6iL exhibited expansion of ΔPE-Oct4-GFP+ cells for at least 8 passages (Figure 5L).

These findings demonstrate that the 6iL condition provides a robust and promising system for cultivating naive hiPSCs and hESCs, supporting their pluripotency. Notably, the basal medium E4 in the 6iL culture system is serum-free, which is of great significance for clinical and pre-clinical applications.

## DISCUSSION

In this study, we demonstrated that the ‘6iL/E4’ culture system enables the derivation and long-term maintenance of ESCs from diverse mammalian species. These findings support the hypothesis that the fundamental mechanisms governing ESC self-renewal are likely conserved across mammals. Deriving ESCs under a shared culture condition provides a unified platform for studying species-specific developmental processes, modeling diseases, and advancing regenerative medicine. For example, while mice and rats have well-established interspecies chimera models, the development of similar tools in other mammals remains technically challenging or unachievable using current approaches.^37–39^ By validating the 6iL ESC culture condition across multiple species, we offer a broadly applicable system to enhance interspecies chimerism and enable ESC-based applications in non-rodent models. The successful derivation of chimera-competent rabbit and bovine ESCs using 6iL represents a significant advance, paving the way for basic and translational research and conservation efforts through advanced reproductive technologies.

Through systematic dissection of ESC self-renewal signaling pathways, several key insights emerged. First, we identified GSK3α-specific inhibition as critical for the derivation of ESCs from non-rodent species. While CHIR, an inhibitor of both GSK3α and GSK3β, promotes self-renewal in mouse and rat ESCs by stabilizing β-catenin,^3,7,40^ it induced substantial differentiation in rabbit and bovine cultures (Figure 2A). In contrast, the WNT pathway inhibitor IWR1 effectively suppressed non-ESC outgrowths (Figure 2A). This observation aligns with recent protocols for culturing human and cynomolgus monkey naive PSCs,^41,42^ which exclude CHIR in favor of WNT inhibitors like IWR1 or XAV939. Additionally, species-specific differences in β-catenin signaling have been reported: in mouse ESCs, β-catenin promotes self-renewal, whereas in human naive ESCs, β-catenin overexpression induces differentiation, and its deletion sustains pluripotency.^18^ Interestingly, low doses of CHIR (1/10 to 1/3 of standard mESC concentrations) can support human naive ESCs, indicating that GSK3 inhibition may still play a role via β-catenin–independent mechanisms.^30,43^ Our recent studies revealed that the effects of CHIR in mESC stemness maintenance are primarily due to GSK3β inhibition, which in turn promotes the nuclear translocation of β-catenin,^40^ whereas the GSK3α-specific inhibitor BRD0705 sustains mESC and mEpiSC self-renewal through a β-catenin–independent mechanism.^19^ Moreover, the combination of BRD0705 and IWR1 sustained the distinct identities of mESCs and mEpiSCs over extended culture periods. Here, we confirmed that the BRD0705/IWR1-based 6iL condition is essential for deriving ESCs from multiple species. Notably, substituting BRD0705 with CHIR during rabbit and bovine ESC derivation led to pronounced differentiation (Figure 1M), highlighting the pivotal role of GSK3α-specific inhibition in establishing stable ESC cultures for nonrodent mammals.

Second, we have identified the critical role of the PDGFR inhibitor CP673451 in the derivation and culture of ESCs across multiple species. Notably, 6iL-ESCs exhibit comparable pluripotency to established naive ESCs in both rodents and humans, suggesting that inhibition of PrE differentiation is critical for efficient epiblast embryonic stem cell derivation. Although further investigation to determine whether PDGF inhibitors can suppress endoderm development is warranted, our observation is supported by previous studies. For example, during human embryonic development, PDGF is expressed in all three blastocyst lineages, epiblast (EPI), extra-embryonic primitive endoderm (PrE), and trophectoderm (TE),^44,45^ while PDGFR expression is prominently observed in the PrE.^46^ Similarly, in mice, PDGF is highly expressed across pre-implantation embryo stages while PDGFR is highly expressed in the PrE.^47,48^ Activation of the PDGF signaling pathway by adding PDGF is crucial for PrE specification during mouse embryonic development and its derivation and expansion in vitro.^49^ Cells captured at the epiblast stage in both humans and mice have been shown to retain pluripotency.^41,50^ Notably, 6iL-ESCs derived from various mammalian species exhibit pluripotency, which strongly suggesting that the inhibition of PrE differentiation is critical for the efficient derivation of ESCs.

Third, we have, for the first time, identified that the LATS1/2 inhibitors TDI/Truli significantly promote the self-renew of rabbit and bovine ESCs. It has been shown that knocking down YAP in mESCs results in reduced stemness, whereas ectopic expression of YAP prevents differentiation and sustains stem cell phenotypes even under differentiation-inducing conditions.^51^ Similarly, overexpression of YAP in hESCs facilitates a transition from the post-implantation primed state to the pre-implantation naive state.^52^ Our findings further extend these studies and highlight the pivotal and conserved role of the YAP signaling pathway in supporting ESC proliferation across multiple mammalian species.

Fourth, it was known that mESCs derived from ICM and cultured in 2iL are typically correspond to the E4.5 developmental stage,^53^ enabling them to remain in a long-term naive state.^7^ However, whether ESCs from non-rodent mammals can be maintained in a long-term naive state remains unknown. We found that the responses to MEK and GSK3 signaling pathways in the establishment of ESCs from rabbit and bovine embryos are completely opposite to those observed in mice, making it challenging to capture their naive states as readily as in mice. This discrepancy is likely attributable to the inherent embryonic characteristics of mice. Notably, mouse embryos undergo diapause,^54^ a hormonally regulated developmental arrest at the E3.5 stage,^55,56^ that allows an extended time window for capturing cells in the naive state for ESC derivation. In contrast, pre-implantation development in higher mammals, such as humans, is continuous, and there is no evidence that these species exhibit embryonic diapause similar to that of mice.^54^ This likely explains why 2i fails to derive ESCs from higher mammals and naive ESCs cannot be established from these species. Notably, we presented a solution for large animals such as bovine in this study-by transiently overexpressing pluripotency genes such as Klf2 and Nanog, bovine ESCs can be artificially reverted to an earlier developmental stage, thereby enhancing their chimera-forming capacity and providing a powerful means to generate genetically engineered animals.

In conclusion, we have comprehensively tested key signaling pathways to capture and maintain mammalian pluripotency, and developed a universal 6iL/E4 culture system for deriving and maintaining ESCs across mammalian species. We confirmed the conservation of the WNT, STAT3, and FGF signaling pathways during the derivation and expansion of ESCs across multiple species, and for the first time, we identified the role of the PDGF and HIPPO signaling pathways in the establishment of these cells, thereby expanding our understanding of ESC self-renewal across species. The established and characterized robust multi species ESC lines hold immense promise for various applications. The universal 6iL ESC culture system provides a powerful gateway to understanding of the signaling pathways required to maintain pluripotent stem cells across species, as well as for the derivation and expansion of a series of mammalian ESCs that have not been previously established.

### Limitations of Study

We thoroughly tested signaling regulators capturing the pluripotency across mammalian species and developed a powerful 6iL culture system; however, the culture system admittedly adds complexity compared to existing approaches, requiring a number of small molecules and growth factors. Further refinements are needed to identify the effective and minimal essential components required for robust ESC maintenance across a broad range of species. Additionally, its performance varies across species, necessitating further optimization to support the derivation and expansion of ESCs from particularly recalcitrant species. Insights from such studies will not only deepen our understanding of pluripotency but also accelerate the development of ESC-based applications in regenerative medicine, agriculture, and biodiversity conservation. Although 6iL enables the derivation of chimera-competent bESCs, germline transmission remained uncharacterized in this study due to pronounced challenges of performing such experiments in a large mammal. Addressing these challenges will unlock the full potential of ESC technology across the mammalian kingdom.

## METHODS

### Key resources table

| REAGENT or RESOURCE | SOURCE | IDENTIFIER |
| --- | --- | --- |
| **Antibodies** |  |  |
| Mouse monoclonal anti-MF 20 | Developmental Studies Hybridoma Bank | CAT#MF 20; RRID: AB_2147781 |
| Rabbit monoclonal anti-FOXA2 | Cell Signaling Technology | CAT#8186; AB_10891055 |
| Rabbit monoclonal anti-TUJ1 | Cell Signaling Technology | CAT#5568; RRID: AB_10694505 |
| Rabbit monoclonal anti-NANOG | Thermo Fisher Scientific | CAT#500-P236-50ug; RRID: AB_2929968 |
| Mouse monoclonal anti-OCT3/4 Antibody (C-10) | Santa Cruz | CAT#sc5279; RRID: AB_628051 |
| Mouse monoclonal anti-OCT3/4 (A-9) | Santa Cruz | CAT#sc365509; RRID: AB_10859218 |
| Rabbit polyclonal anti-NANOS3 | GeneTex | CAT#GTX85389; RRID: AB_10726267 |
| Rabbit monoclonal anti-YAP | Cell Signaling Technology | CAT#14074; RRID: AB_2650491 |
| Goat Polyclonal anti-GFP | Rockland | CAT#600-101-215; RRID: AB_11181883 |
| Donkey anti-Goat IgG (H+L) Cross-Adsorbed Secondary Antibody, Alexa Fluor™ 488 | Invitrogen | CAT#A-11055; RRID: AB_2534102 |
| Mouse monoclonal anti-TFAP2C | Santa Cruz | CAT#sc-12762; RRID: AB_667770 |
| Goat polyclonal anti-mouse IgG-H&L (Cy5) | abcam | CAT#ab6563; RRID: AB_955068 |
| Goat polyclonal anti-mouse IgG-H&L (Cy5) | Invitrogen | CAT#A10524; RRID: AB_10562712 |
| Goat polyclonal anti-rabbit IgG, Alexa Fluor™ Plus 488 | Invitrogen | CAT#A32731; RRID: AB_2633280 |
| Donkey polyclonal anti-Rabbit IgG (H+L), Alexa Fluor™ 555 | Invitrogen | CAT#A31572; RRID: AB_162543 |
| Goat polyclonal anti-PRDM1 | LS Bio | CAT#LS-C112748-100; RRID: AB_10710210 |
| Mouse monoclonal anti-GATA4 | Santa Cruz | CAT#sc25310; RRID: AB_627667 |
| Mouse monoclonal anti-CTNT | Thermo Fisher Scientific | CAT#MA5-12960; RRID: AB_11000742 |
| Rabbit monoclonal anti-OCT4 | Thermo Fisher Scientific | CAT#MA5-14845; RRID: AB_10979606 |
| Chemicals, peptides, and recombinant proteins |  |  |
| IWR1 | Selleckchem | CAT#S7086 |
| BRD0705 | Cayman | CAT#37314 |
| CP673451 | Selleckchem | CAT#S1536 |
| GDC0879 | Selleckchem | CAT#S1104 |
| Recombinant Human LIF | Gibco | CAT#300-05-5UG |
| Recombinant Bovine LIF | Kingfisher Biotech | CAT#RP0997B |
| 828 | WuXi AppTec | N/A; Company-synthesized. |
| SKL2001 | Selleckchem | CAT#S8320 |
| TRULI | Selleckchem | CAT#E1061 |
| TDI-011536 | Selleckchem | CAT#E1314 |
| SU5402 | Selleckchem | CAT#S7667 |
| Axitinib | Selleckchem | CAT#S1005 |
| Futibatinib | Selleckchem | CAT#S8848 |
| PD184352 | Selleckchem | CAT#S1020 |
| Vx11e | Selleckchem | CAT#S7709 |
| G06983 | Selleckchem | CAT#S2911 |
| JNK-IN-8 | Selleckchem | CAT#S4901 |
| CHIR-99021 | Selleckchem | CAT#S1263 |
| PD0325901 | Selleckchem | CAT#S1036 |
| N2 Supplement | Gibco | CAT#17502048 |
| B27 Supplement | Gibco | CAT#17504044 |
| Neurobasal medium | Gibco | CAT#21103049 |
| DMEM/F12 medium | Gibco | CAT#11320033 |
| StemSpan™ SFEM II | Stemcell technologies | CAT# 09655 |
| Gelatin | Sigma-Aldrich | CAT#G1890-500G |
| L-Glutamin | Gibco | CAT#25030081 |
| MEM NEAA | Gibco | CAT#11140050 |
| Sodium Pyruvate | Gibco | CAT#11360070 |
| IMDM | Gibco | CAT#12440053 |
| DMEM | Gibco | CAT#11965092 |
| Human holo-transferrin | Sigma-Aldrich | CAT#T0665 |
| BSA | Sigma-Aldrich | CAT#A8806-5G |
| Recombinant Human Insulin | Sigma-Aldrich | CAT#91077C |
| Doxycyline | Sigma-Aldrich | CAT#D5207-1G |
| Sodium selenite | Sigma-Aldrich | CAT#S5261 |
| Bovine holo-transferrin | Sigma-Aldrich | CAT#T1283-50MG |
| Corticosterone | Sigma-Aldrich | CAT#C2505 |
| D+-galactose | Sigma-Aldrich | CAT#G0625 |
| DL-alpha tocopherol acetate | Sigma-Aldrich | CAT#T3001-10G |
| Progesterone | Sigma-Aldrich | CAT#P8783 |
| Linolenic acid | Sigma-Aldrich | CAT#L2376 |
| Glutathione reduced | Sigma-Aldrich | CAT#G6013 |
| Biotin | Sigma-Aldrich | CAT#B4639 |
| T3 | Sigma-Aldrich | CAT#T6397 |
| Catalase | Sigma-Aldrich | CAT#C40-100MG |
| Sodium selenite | Sigma-Aldrich | CAT#S9133 |
| Putrescine | Sigma-Aldrich | CAT#P5780 |
| Vit E | Sigma-Aldrich | CAT#T3251 |
| Linoleic acid | Sigma-Aldrich | CAT#L1012 |
| BO-IVM medium | IVF Bioscience | CAT#71001 |
| BO-IVF medium | IVF Bioscience | CAT#71004 |
| BO-IVC medium | IVF Bioscience | CAT#71005 |
| **Critical commercial assays** |  |  |
| Alkaline Phosphatase Kit | Vector laboratories | CAT#SK-5300 |
| RNeasy Mini Kit | Qiagen | CAT#74104 |
| iTaq™ Universal SYBR® Green Supermix | Bio-Rad | CAT#1725124 |
| **Deposited data** |  |  |
| 6iL-mESC\| RNA-seq datasets | This paper | GEO: GSE295390<br>https://www.ncbi.nlm.nih.gov/geo/query/acc.cgi?acc=GSE295390 |
| 6iL-hiPSC\| RNA-seq datasets | This paper | GEO: GSE295412<br>https://www.ncbi.nlm.nih.gov/geo/query/acc.cgi?acc=GSE295412 |
| **Experimental models: Cell lines** |  |  |
| Mouse: mESCs | This paper | N/A |
| Mouse: 6iL-mESCs | This paper | N/A |
| Mouse: GFP 6iL-mESCs | This paper | N/A |
| Rat: DAC8 rESCs | Li et al.<sup>4</sup> | N/A |
| Bovine: 6iL-bESCs | This paper | N/A |
| Bovine: GFP 6iL-bESCs | This paper | N/A |
| Bovine: GFP-KLF2/NANOG/M2RTta 6iL-bESCs | This paper | N/A |
| Rabbit: 6iL/TDI-rabESCs | This paper | N/A |
| Rabbit: GFP 6iL/TDI-rabESCs | This paper | N/A |
| Human: 6iL-hiPSCs | This paper | N/A |
| Human: ΔPE-Oct4-GFP WIBR3 hESCs | Theunissen et al.<sup>43</sup> | N/A |
| Human: KLF2/NANOG/M2rtTA ΔPE-Oct4-GFP WIBR3 hESCs | This paper | N/A |
| **Experimental models: Organisms/strains** |  |  |
| Mouse/B6D2F1 | Charles river | 099 |
| Mouse/C57BL/6J | Charles river | 027 |
| Rabbit/New Zealand White | Inotiv | 221 |
| Bovine/Angus | University of Florida/IFAS Beef Teaching Unit | N/A |
| Bovine embryos/IVF | University of Florida/Animal | N/A |
|  | science department |  |
| Oligonucleotides |  |  |
| Primers for qRT-PCR | See Table S1 | N/A |
| Recombinant DNA |  |  |
| FUW-tetO-loxP-hKLF2 | Addgene | RRID: Addgene_60850 |
| FUW-tetO-loxP-hNANOG | Addgene | RRID: Addgene_60849 |
| FUW-M2rtTA | Addgene | RRID: Addgene_20342 |
| pCXLE-hOCT3/4-shp53-F | Addgene | RRID: Addgene_27077 |
| pCXLE-hUL | Addgene | RRID: Addgene_27080 |
| pCXLE-hSK | Addgene | RRID: Addgene_27078 |
| Software and algorithms |  |  |
| Fiji-ImageJ | National Institutes of Health | https://imagej.net/Fiji |
| GraphPad Prism 9 | GraphPad | https://www.graphpad.com/scientific-software/prism/ |
| Snapgene | Snapgene | https://www.snapgene.com |
| STAR | Dobin et al.<sup>57</sup> | https://code.google.com/archive/p/rna-star/;<br>RRID: SCR_004463 |
| DESeq2 v1.40.2 | Love et al.<sup>58</sup> | http://www.bioconductor.org/packages/release/bioc/html/DESeq2.html. |
| RSEM | Li et al.<sup>59</sup> | http://github.com/deweylab/RSEM |
| Other |  |  |
| Human pluripotent stem cell \| RNA-seq datasets | Irie et al.<sup>35</sup> | GEO: GSE60138<br>(https://www.ncbi.nlm.nih.gov/geo/query/acc.cgi?acc=GSE60138) |
| Human pluripotent stem cell \| RNA-seq datasets | Takashima et al.<sup>30</sup> | ArrayExpress: E-MTAB-2856<br>(https://www.ebi.ac.uk/biostudies/ArrayExpress/studies/E-MTAB-2856?query=E-MTAB-2856) |
| Human pluripotent stem cell \| RNA-seq datasets | Guo et al.<sup>36</sup> | ArrayExpress: E-MTAB-4462<br>(https://www.ebi.ac.uk/biostudies/arrayexpress/studies/E-MTAB-4462) |
| Mouse pluripotent stem cell \| RNA-seq datasets | Kinoshita et al.<sup>50</sup> | GEO: GSE131556<br>(https://www.ncbi.nlm.nih.gov/geo/query/acc.cgi?acc=GSE131556) |

## RESOURCE AVAILABILITY

### Lead Contact

For further inquiries and requests for resources and reagents, please contact the Lead Contact, Qilong Ying.

### Materials Availability

All regents and stable cell lines associated with this manuscript can be obtained from the Lead Contact, provided that a Materials Transfer Agreement from the University of Southern California is completed.

## EXPERIMENTAL MODEL AND SUBJECT DETAILS

### Mice

Adult female mice were used in these experiments. B6D2F1 mice were used for cell line derivation, C57BL/6J mice provided host embryos for chimera generation, and ARC mice served as embryo transfer recipients. Embryonic stem cell injections, blastocyst transplantation, and post-transfer mouse culture were conducted at the Irvine Transgenic Mouse Facility, University of California, Irvine, under IACUC protocol #AUP-22-126. Mouse embryo collection for mESC derivation was performed at the University of Southern California’s Department of Animal Resources and all animal experiments were performed according to the investigator’s protocols approved by the University of Southern California Institutional Animal Care and Use Committee. The project was approved by the Animal Welfare and Ethical Review Bodies of both institutions.

### Bovine

Non-lactating, 3-year-old crossbreed (Bos taurus x Bos indicus) cows were used as recipient cows for chimera experiments. The animal experiments were conducted under animal use protocols (202300000191) approved by the Institutional Animal Care and Use Committee of the University of Florida. All cows were housed in open pasture, and under constant care of the farm staff.

### Rabbit

New Zealand White (NZW) rabbits were used in this study. The animal maintenance, care, and use procedures were reviewed and approved by the Institutional Animal Care and Use Committee (IACUC, protocol #PRO00011844) of the University of Michigan. All procedures were carried out in accordance with the approved guidelines.

### Harvesting and culture of mouse embryos

Female B6D2F1 mice (8–10 weeks old) were induced to superovulate via an intraperitoneal injection of 5 IU PMSG (Prospec), followed 48 hours later by an intraperitoneal injection of 5 IU hCG. After mating with C57BL/6 males, embryos ranging from the 2-cell stage were collected at embryonic day 2.75 (E2.75) from the oviducts and uterine horns using KSOM; the detection of a vaginal plug was designated as embryonic day 0.5 (E0.5). The embryos were then cultured at 37℃ in atmosphere containing 5% (v/v) CO2. At E3.5, blastocysts were flushed from the uterine horns.

### Bovine in vitro embryo production

Germinal vesicle (GV) stage oocytes were collected as cumulus–oocyte complexes (COCs) aspirated from slaughterhouse ovaries. In vitro maturation was performed in BO-IVM medium (IVF Bioscience, Falmouth, UK) at 38.5°C with 6% CO₂ for 22–23 hours to obtain metaphase II (MII) oocytes. Cryopreserved semen from a Holstein bull with proven fertility was prepared in BO-SemenPrep medium (IVF Bioscience, Falmouth, UK) and added to drops containing COCs at a final concentration of 2 × 10⁶ spermatozoa/ml for in vitro fertilization. Gametes were co-incubated at 38.5°C and 6% CO₂. After 10 hours (for microinjection experiments) or 16 hours (for non-microinjection experiments) in BO-IVF medium (IVF Biosciences, Falmouth, UK), IVF embryos were denuded of cumulus cells by vortexing for 5 minutes in BO-Wash medium (IVF Bioscience, Falmouth, UK) and then cultured for up to blastocyst stage on 7.5 day in BO-IVC medium (IVF Biosciences, Falmouth, UK) at 38.5°C, 6% CO₂, and 6% O₂. Embryos at various developmental stages were evaluated under light microscopy according to the International Embryo Technology Society’s grading standards.

### Rabbit embryos collection and culture

Superovulation, embryo collection and culture were conducted as previously described.^60^ Briefly, adult NZW female rabbits were superovulated with follicle-stimulating hormone (FSH, Folltropin-V, Bioniche Life Sciences, Canada) and human chorionic gonadotropin (hCG, Chorulon, Intervet, Millsboro, DE) to induce ovulation, followed by breeding with male rabbits. Eighteen hours post breeding, the zygote stage embryos were collected, and cultured in the embryo culture medium in vitro to blastocyst stage. The embryo culture medium is composed of 10% fetal bovine serum (FBS, 10438-026, Thermofisher), MEM non-essential amino acids (M7145, Thermofisher), BME amino acid solution (B6766, Milliporesigma, Burlington, MA, USA), 2 mM L-glutamine (25030081, Thermofisher, Waltham, MA, USA), 0.4 mM sodium pyruvate (11360070, Thermofisher) in Earl’s Balanced Salts (E2888, Milliporesigma).

## METHOD DETAILS

### Derivation and culture of 6iL-mESCs

E3.5 mouse blastocysts were briefly exposed to acidic Tyrode’s solution to remove their zona pellucida (ZP). Following this, the de-zonated embryos were plated on MEFs using the 6iL medium to derive mESCs (E4 medium supplemented with human LIF (20 ng/mL, Peprotech), IWR1 (2.5µM, Selleck), BRD0705 (8µM, Cayman), CP673451 (1µM, Canyon), GDC0879 (1µM, Selleck), 828 (5µM, WuXi AppTec), SKL2001 (10µM, Selleck)). Additionally, during the culture process, Go6983 (1 µM, Selleck) can be selectively added to make the stem cell clones more compact. After being cultured for 4–6 days, blastocyst outgrowths were dissociated using 0.025% trypsin and transferred onto freshly prepared MEFs for further cultivation. The cells were cultured at 37℃ under 5% CO2. E4 medium: the 1:1 mixture of DMEM/F12 and Neurobasal medium, supplemented with insulin (4 μg/ml, Sigma), human holo-transferrin (22 μg/ml, Sigma), BSA (1 mg/ml, Sigma), sodium selenite (12.5 ng/ml, Sigma), and L-glutamine (2 mM, ThermoFisher).

### Derivation and culture of 6iL-rabESCs

Zona pellucida–removed rabbit morula embryos were placed on MEFs and initially cultured in E4 medium supplemented with human LIF (20 ng/mL, Peprotech), IWR1 (2.5 µM, Selleck), BRD0705 (8 µM, Cayman), CP673451 (1 µM, Canyon), GDC0879 (1 µM, Selleck), 828 (5 µM, WuXi AppTec), SKL2001 (10 µM, Selleck), and TDI-011536 (100nM, Selleck) for 4–6 days. The outgrowths were then dissociated using 0.025% trypsin and transferred onto freshly prepared MEFs for further cultivation. The cells were maintained at 38.5°C under 5% CO₂.

### Derivation and culture of 6iL-bESCs

ICMs isolated from bovine blastocysts were placed on MEFs and initially cultured in E4 medium (with additional 50µg/ml bovine transferrin (Sigma, T1283)) supplemented with bovine LIF (20 ng/mL, Kingfisher Biotech), IWR1 (2.5 µM, Selleck), BRD0705 (8 µM, Cayman), CP673451 (1 µM, Canyon), GDC0879 (1 µM, Selleck), 828 (5 µM, WuXi AppTec), and SKL2001 (10 µM, Selleck) can also be added for 4–6 days. SKL2001 can be replaced with TRULI (2 µM, Selleck) or TDI-011536 (100nM, Selleck). The outgrowths were then dissociated using 0.025% trypsin and transferred onto freshly prepared MEFs for further cultivation. bESC clones were picked and then dissociated for passaging. The cells were maintained at 38.5°C under 5% CO₂.

### Derivation and culture of human iPSC using 6iL

Cells obtained from centrifuged umbilical human cord blood were electroporated with the plasmids pCXLE-hOCT3/4-shp53, pCXLE-hSK, and pCXLE-hUL.^34^ After culturing the human cord blood cells in StemSpan™ SFEM II medium (STEMCELL technologies) for two days, the medium was replaced with E4 medium supplemented with LIF (20 ng/mL, Peprotech), IWR1 (2.5 µM, Selleck), BRD0705 (8 µM, Cayman), CP673451 (1 µM, Canyon), GDC0879 (1 µM, Selleck), 828 (5 µM, WuXi AppTec), SKL2001 (10 µM, Selleck), and the cells were cultured on MEFs. SKL2001 can be replaced with TRULI (2 µM, Selleck) or TDI-011536 (100nM, Selleck). Additionally, during the culture process, Go6983 (1 µM, Selleck) can be selectively added to make the stem cell clones more compact. After approximately 10–12 days, clones were picked and passaged onto feeder-coated plates using 0.025% trypsin digestion. The cells were maintained at 37°C under 5% CO₂.

### 6iL-rESC culture

The DAC8 rat ESC line was maintained on MEF plates pre-coated with 0.1% gelatin in E4 medium supplemented with 6iL. The cultures were incubated at 37°C with 5% CO₂, and the medium was changed daily. For passaging, cells were dissociated into single cells using 0.025% trypsin.

### 6iL-Naïve human ESC culture

OCT4-ΔPE-GFP-WIBR3 hESCs were infected with lentiviruses (FUW-tetO-lox-hKLF2, FUW-tetO-lox-hNANOG, and M2rtTA). The cells were cultured on plates with an MEF feeder layer in E4 medium supplemented with CHIR/PD03/LIF, with DOX added for selection over three passages. GFP-positive single clones were then selected and cultured on MEF plates pre-coated with 0.1% gelatin in the presence of human LIF (20 ng/mL, PeproTech), IWR1 (2.5 µM, Selleck), BRD0705 (8 µM, Cayman), CP673451 (1 µM, Canyon), GDC0879 (1 µM, Selleck), 828 (5 µM, WuXi AppTec), SKL2001 (10 µM, Selleck). Additionally, during the culture process, Go6983 (1 µM, Selleck) can be selectively added to improve cell condition. The cultures were incubated at 37°C with 5% CO₂, and the medium was changed daily. For passaging, cells were dissociated into single cells using 0.025% trypsin.

### Alkaline phosphatase (AP) staining

The AP substrate solution (Vector Laboratories) was prepared following the manufacturer’s instructions. Cells were incubated with the AP substrate at room temperature for 20–30 minutes in the dark. After incubation, they were fixed with 4% (w/v) paraformaldehyde (PFA) at room temperature for 1 hours.

### qRT-PCR Analysis

Total RNA was extracted using the RNeasy Mini Kit (Qiagen) according to the manufacturer’s protocol. cDNA was synthesized using the iScript cDNA Synthesis Kit (Bio-Rad). Quantitative real-time PCR was performed with the iTaq Universal SYBR® Green Supermix (Bio-Rad) on a Viia 7 real-time PCR system. Gene expression levels were normalized to Gapdh.

### Immunofluorescence

Cells were fixed in a 4% PFA solution for 15 minutes at room temperature, followed by three PBS washes. Next, the cells were blocked with 5% BSA in PBS containing 0.3% Triton X-100 for 1 hour. Primary and secondary antibodies—diluted in 1% BSA in PBS with 0.3% Triton X-100—were subsequently applied for either 1 hour at room temperature or overnight at 4°C. Details of the antibodies used can be found in the Key Resources Table.

### EBs formation and differentiation

EBs were formed using AggreWell 400 plates (Stem Cell Technologies) in accordance with the manufacturer’s protocol. EBs from different species were cultured for 2–4 days. For cardiomyocyte differentiation, the resulting EBs were plated onto gelatin-coated dishes and cultured in either IMDM/10% FBS or GMEM/10% FBS medium. For neural differentiation, EBs were plated onto gelatin-coated dishes and maintained in N2B27 medium.

### Primordial germ cell like cell (PGC-LC) induction

EBs were formed using AggreWell 400 plates (Stem Cell Technologies). The basic steps for PGC-LC induction and the culture medium recipe were carried out according to a published protocol.^31,61,62^ Rat and bovine are first cultured overnight in N2B27 medium containing Activin A (20 ng/ml, Stem Cell Technologies) and bFGF (20 ng/ml, Peprotech). 6iL-hiPSCs can be directly placed into PGC induction medium. EB formation occurs in the PGC-LC induction medium: GK15 medium containing GMEM (15% (v/v) KnockOut serum replacement (ThermoFisher), NEAA (0.1 mM, ThermoFisher), sodium pyruvate (0.1mM, ThermoFisher), b-mercaptoethanol (0.1 mM, Sigma), Glutamax (2 mM, ThermoFisher)), supplemented with BMP4 (200 ng/mL, GIBCO), LIF (1000 U/mL; Peprotech), SCF (100 ng/mL; R&D) and EGF (50 ng/mL; Peprotech).

### Lentiviral infection

Lentiviruses pseudotyped with VSVG and PSPAX were produced in HEK-293 cells following established protocols. In brief, the culture medium was replaced 12 hours after transfection, and the virus-containing supernatant was harvested between 48 and 72 hours post-transfection. The collected supernatant was then filtered through a 0.45 µm filter. Finally, the viral supernatants (FUW-tetO-loxP-hKLF2, FUW-tetO-loxP-hNANOG, and FUW-M2rtTA) were added to the 6iL-bESCs or hESCs.Two rounds of infection were carried out over a 24-hour period, each in the presence of 2 μg/ml polybrene.

### RNA-seq library preparation and data analysis

Total RNA from individual cell lines was extracted using RNeasy Micro Kit (Qiagen). The RNA-seq libraries were generated using the Smart-seq2 v4 kit and Nextera XT DNA Library Preparation Kit (Illumina), and multiplexed by Nextera XT Indexes (Illumina) following the manufacturer’s instructions. The concentration of sequencing libraries was determined using Qubit dsDNA HS Assay Kit (Life Technologies) and KAPA Library Quantification Kits (KAPA Biosystems). The size of sequencing libraries was determined using the Agilent D5000 ScreenTape with Tapestation 4200 system (Agilent). Pooled indexed libraries were then sequenced on the Illumina NovaSeq platform with 150-bp pair-end reads.

Multiplexed sequencing reads that passed filters were trimmed to remove low-quality reads and adaptors by Trim Galore (version 0.6.7) (-q 25 –length 20 –max_n 3 –stringency 3). The quality of reads after filtering was assessed by FastQC, followed by alignment to the bovine genome (ARS-UCD1.3) by HISAT2 (version 2.2.1) with default parameters. The output SAM files were converted to BAM files and sorted using SAMtools6 (version 1.14). Read counts of all samples were quantified using featureCounts (version 2.0.1) with the reference genome. Principal component analysis and cluster analysis were performed with R (a free software environment for statistical computing and graphics). Differentially expressed genes (DEGs) were identified using edgeR in R. Genes were considered differentially expressed when they provided a false discovery rate of <0.05 and fold change >2. ClusterProfiler was used to reveal the Gene Ontology and KEGG pathways in R.

### Bovine in vivo chimera assay

Five to ten GFP+ bESCs were injected gently into the morula or early blastocyst stage bovine embryos using a piezo-assisted micromanipulator attached to an inverted microscope (Olympus). The injected embryos were cultured in BO-IVC and bESCs mixture medium (75%:25%) at 38.5 °C, 6% CO_2_, and 6% O_2_ for extra 8 hours. The injected blastocysts with clear GFP signaling were then transferred into non-lactating, 3-year-old crossbreed recipient cows (n = 10). Recipient cows were synchronized with a standard 7-day controlled internal drug release (CIDR, Zoetis) protocol, following with one IM dose of ovulation-inducing gonadotrophin releasing hormone (Fentagyl, Merk Aninmal Health). At day 7, CIDR was removed, and one dose of Prostaglandin (Lutalyse, Zoetis) was administered. On day 9, one dose of Fertagyl was administered to stimulate ovulation. Subsequently the transfer was made 7 days later. At day 30 after transplantation, pregnancy was diagnosed by ultrasonography. The recipient cows were slaughtered at the University of Florida Meat Processing Center and the reproductive tracts were harvested to collect day 40 fetus to analysis chimeric competence.

### Cryosection and immunofluorescence analysis of bovine fetus

For immunofluorescence staining of cryo-sections of bovine chimeric fetus, fetuses were collected from uterus and washed with PBS for three times, then they were immersed in 4% PFA overnight at 4°C. After washing with PBS, they were dehydrated sequentially in 10%, 20%, 30% sucrose, OCT:30% sucrose (1:1), 4 hours for each step. Next, fetuses were embedded in Tissue plus O.C.T. compound (Fisher, 4585) and hold on dry ice for quick freezing. The frozen OCT blocks were sectioned by CRYOSTAR NX50 (ThermoFisher), at 10 μm each section. Sections were permeabilized with 1%Triton X-100 in PBS for 30 min and then rinsed with wash buffer. Samples were then transferred to blocking buffer (0.1% Triton X-100, 1% BSA, 0.1 M glycine, 10% donkey serum) for 2 hours at room temperature. Subsequently, the sections were incubated with the primary antibodies overnight at 4°C. The primary antibodies used in this experiment is anti-GFP (Rockland, 600-101-215). For secondary antibody incubation, the cells were incubated with Fluor 488-conjugated secondary antibodies for 1 hour at room temperature. Followed by DAPI staining (Invitrogen, D1306) for 15 min. The images were taken with a fluorescence confocal microscope (Leica).

### Rabbit embryonic stem cells injection and embryo transfer

The ESC injection to 8-cell stage rabbit embryos was carried out as previously described.^60^ Briefly, the rabbit ESCs were trypsinized to single cells before injection. Ten ESCs were injected into each 8-cell stage rabbit embryo. The ESC-injected embryos were either cultured in vitro to evaluate the ESC contribution to the ICM (ICM) at the blastocyst stage embryos or transferred to pseudo-pregnant recipient animals.

### Karyotyping

ESCs were treated with 100 ng/mL colcemid (15212012, Thermofisher) for 2 hours to arrest cell cycle to the metaphase, then trypsinized to single cells. After that, cells were treated with 0.075 M Potassium Chloride (10575090, Thermofisher) for 6 minutes, and then fixed with 25% acetic acid (320099, Milliporesigma) in methanol (34860, Milliporesigma) for 10 minutes at room temperature. After 3 times washing of the cells with 25% acetic acid in methanol, cells were dropped on a glass slide to form chromosome spreads, and the spreads were stained by Giemsa stain (10092-013, Thermofisher). The spreads were randomly pictured under the microscope (BZ800, Keyence, Itasca, IL, US) and manually counted for chromosome number.

### Quantification and statistical analysis RNA-sequencing

Multiplexed sequencing reads that passed filters were trimmed to remove low-quality reads and adaptors by Trim Galore (version 0.6.7). The quality of reads after filtering was assessed by FastQC, followed by alignment to the bovine genome (ARS-UCD1.3) by HISAT2 (version 2.2.1) with default parameters. The output SAM files were converted to BAM files and sorted using SAMtools6 (version 1.14). Read counts of all samples were quantified using featureCounts (version 2.0.1) with the reference genome. Principal component analysis and cluster analysis were performed with R (a free software environment for statistical computing and graphics). Differentially expressed genes (DEGs) were identified using edgeR in R. Genes were considered differentially expressed when they provided a false discovery rate of <0.05 and fold change >2. ClusterProfiler was used to reveal the Gene Ontology and KEGG pathways in R.

### Statistical Analysis

All data were presented as means ± SEM. Experiments were repeated at least three times. Student’s t test (two-tailed) was used to evaluate the statistical significance, and the error bar represents the SEM of three independent experiments. P < 0.05 was taken to indicate statistical significance. *P < 0.05, **P < 0.01 and ***P < 0.001.

## ACKNOWLEDGMENTS

We thank the members of the Ying lab for their technical support. This work was supported by NIH/NIGMS R01 GM129305, NIH Small Business Technology Transfer grant #1R41OD023245, the Chen Yong Foundation of the Zhongmei Group, the Xia Research Fund, and the Wu & Jian Research Fund. Research in the Z.J. laboratory is supported by the NIH Eunice Kennedy Shriver National Institute of Child Health and Human Development Grant (1R01HD102533).

## AUTHOR CONTRIBUTIONS

Conceptualization: D.W. Z.J., J.X. and Q.-L.Y. Experimental Design and Execution: D.W. designed the experiments and performed the majority of the work. D.W. derived 6iL-ESC lines from mouse and bovine blastocysts and from rabbit morulae, and generated 6iL human iPSCs from cord-blood cells. Bovine ESC Work: H.M., and Y.W. produced bovine embryos; H.M., G.S., R.I., and O.O. performed bovine embryo microinjection experiments, chimera assays. Rabbit ESC Work: L.-K.T., Z.W., D.Y., X.K., X.X., and J.Z. produced rabbit embryos and performed rabbit microinjection experiments. RNA-seq and Bioinformatics: L.T., X.W., and G.H. performed bulk RNA sequencing and bioinformatic analyses. Mouse Microinjection: S.W. and K.S. performed mouse microinjection experiments. Compound Provision: D.E. and B.V.H. provided a series of STAT3 activators, including compound 828. qRT-PCR Assistance: K.Y. and B.Z. assisted D.W. with qRT-PCR experiments. Rabbit ESC Project: X.T. initiated and participated in the rabbit ESC project. Human PSC Project: K.Y., L.M., and R.P. contributed to the human PSC project. Supervision: Q.-L.Y., Z.J., and J.X. Funding acquisition: Q.-L.Y. Z.J., and J.X. Resources: Q.-L.Y., Z.J., J.X., and Y.E.C. Manuscript writing: D.W. Q.-L.Y. Z.J., and J.X. wrote the manuscript with input from all authors.

## Supplemental Information

**Figure S1.**
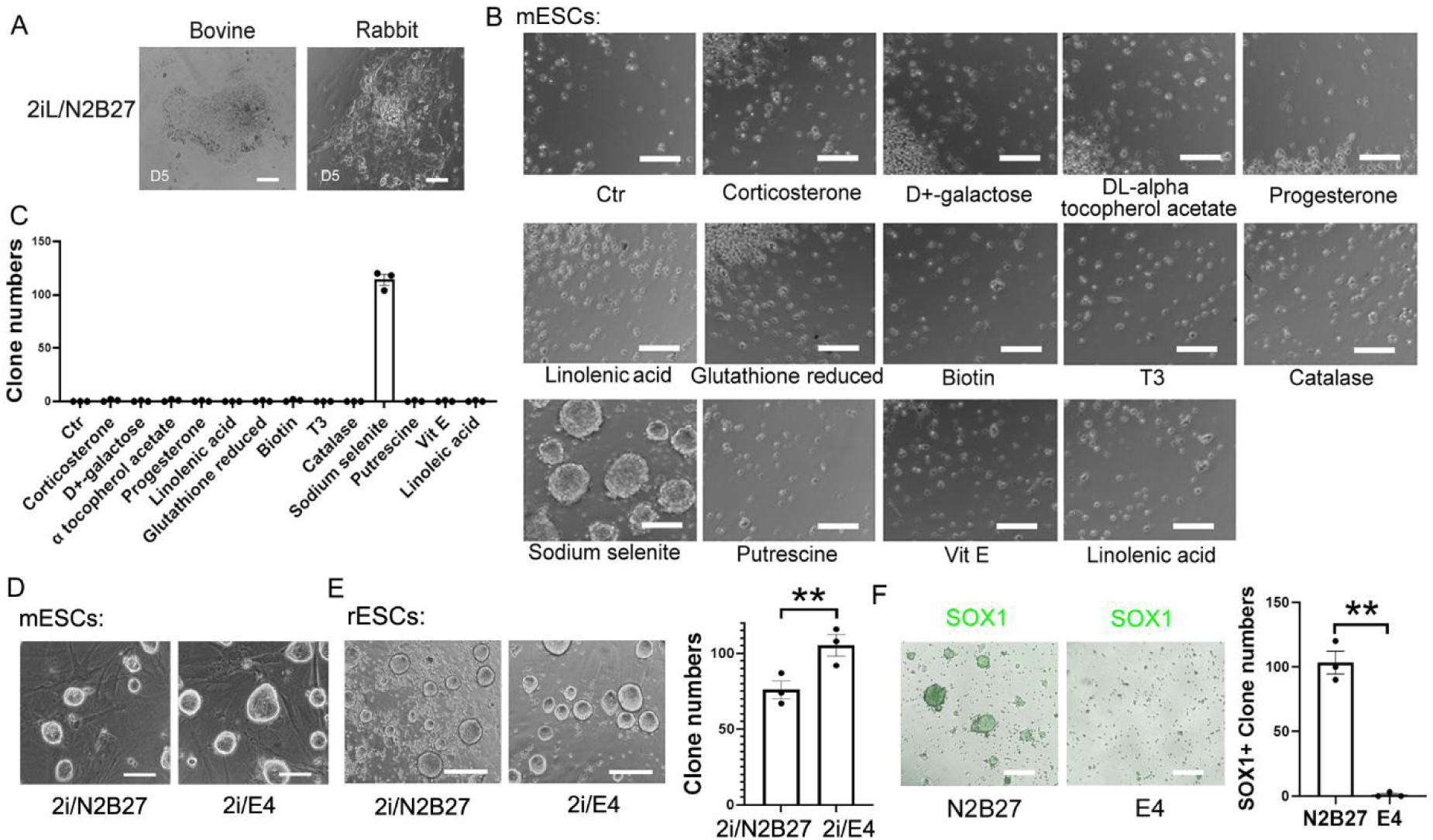
Optimization of N2B27 Composition to Promote ESC self-renewal. (A) Phase-contrast images of bovine and rabbit blastocysts cultured in 2i/N2B27 for 5 days. (B) Phase-contrast images of mESCs cultured for 3 passages on 0.1% gelatin-coated plates in DMEM-F12/Neurobasal medium supplemented with 2i, insulin (4 µg/mL), Tf (20 µg/mL), BSA (1000 µg/mL), and individual B27 components. Scale bar = 200 µm. (C) The bar graph represents the quantitative results of the number of clones in Figure B. (D) Representative phase-contrast images of mESCs cultured for 4 passages in N2B27 or E4 medium supplemented with 2i. Scale bar, 100 µm. (E) Representative phase-contrast images of rESCs cultured for 4 passages in 2i/N2B27 or 2i/E4. Scale bar, 200 µm. The bar graph represents the quantitative results of the number of clones. (F) Representative fluorescence images of Sox1-GFP+ neural stem cells derived from mESCs cultured alone in N2B27 or E4 medium on day 5 of differentiation. Scale bar = 200 µm. Bar chart: quantification of Sox1-GFP+ colonies.

**Figure S2.**
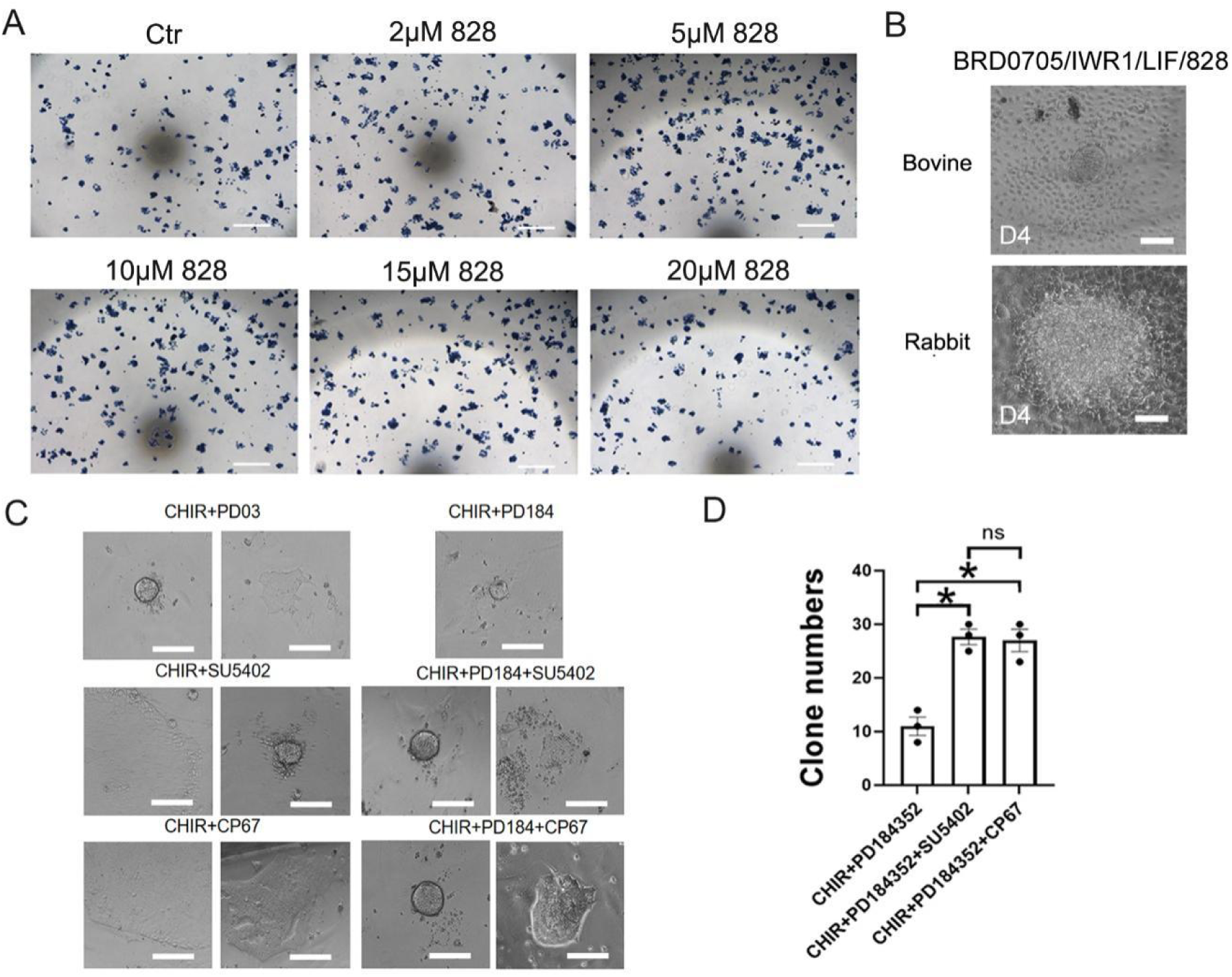
Optimization of Culture Conditions in Rabbit and Bovine ESC Derivation. (A) Alkaline phosphatase (AP) staining of colonies formed by mESCs treated with different concentrations of 828 (0, 2, 5, 10, 15, and 20 μM). Scale bars, 200 μm. (B) Morphology of rabbit and bovine ICMs cultured for 4 days under BRD0705/IWR1/LIF/828 conditions with specific signaling pathway inhibitors. Scale bars, 100 μm. (C) Representative phase-contrast images of rESC cultured with CHIR99021 in combination with different inhibitors: PD03, PD184, SU5402, and CP67. Scale bars, 100 μm. (D) Quantification of clone numbers in figure E. Data are presented as mean ± SEM. Statistical significance is indicated (*P < 0.05, **P < 0.01, ns = not significant).

**Figure S3.**
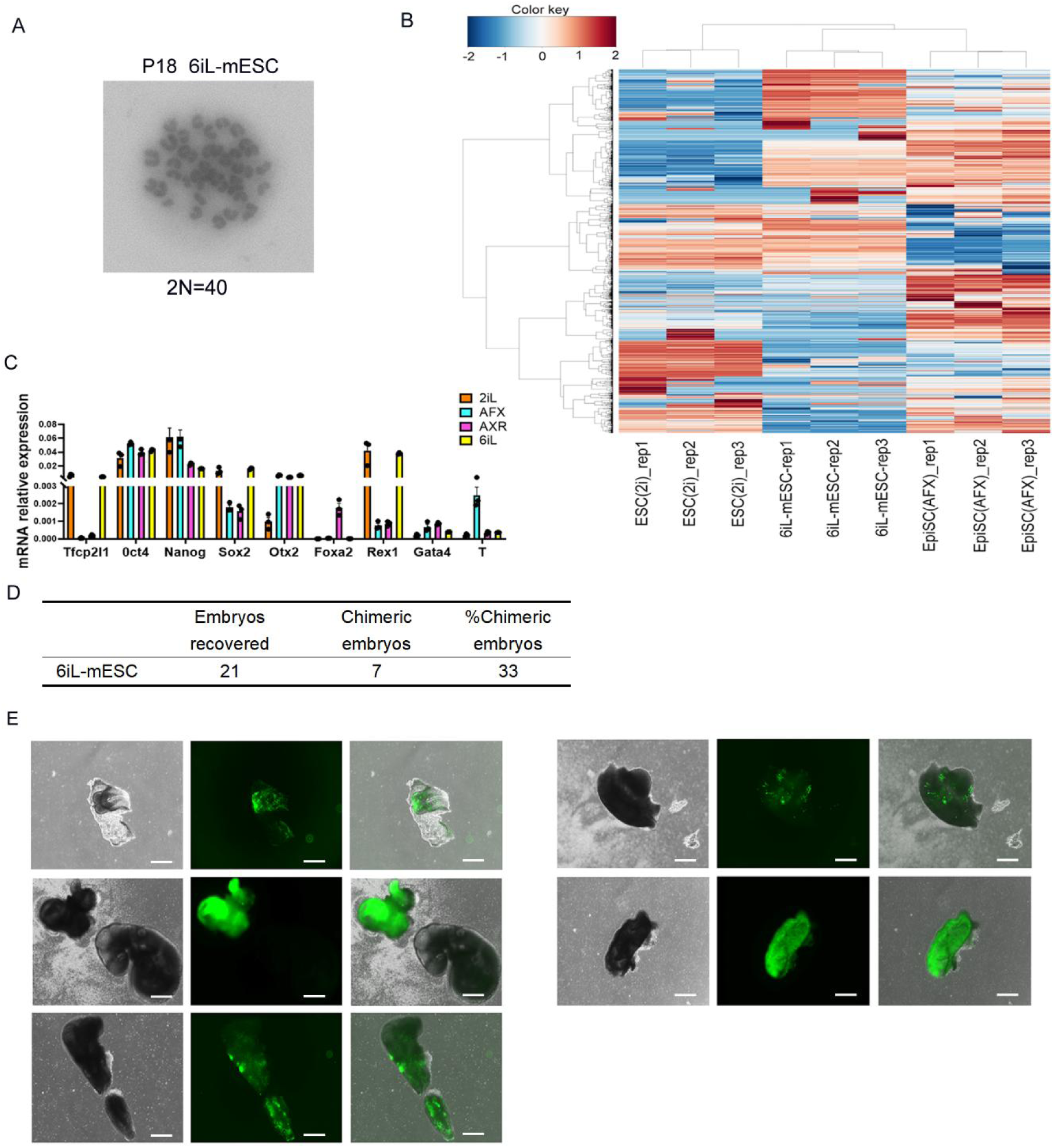
Pluripotency Characterization and Chimeric Contribution of mESCs Derived under 6iL. (A) Representative images showing P18 6iL-mESC karyotyping results. (B) Heatmap showing the global gene expression profile of 6iL-mESCs compared to other ESC lines (2iL and AFX). The hierarchical clustering reveals distinct expression patterns among different culture conditions. (C) qRT-PCR analysis of marker gene expression in Naïve mESC (2iL), primed EpiSC (AFX: Activin A/bFGF/XAV939), formative cell (AloXR: Activin A/XAV939/BMS493) and 6iL ESC. Data are presented as mean ± SEM. (D) Summary table of chimeric contribution efficiency. (E) Representative images of chimeric embryos generated by injecting GFP-labeled 6iL-mESCs into WT blastocyst. Bright-field, fluorescence, and merged images show the distribution of GFP-positive cells in chimeric embryos at different developmental stages. Scale bars, 500 μm.

**Figure S4.**
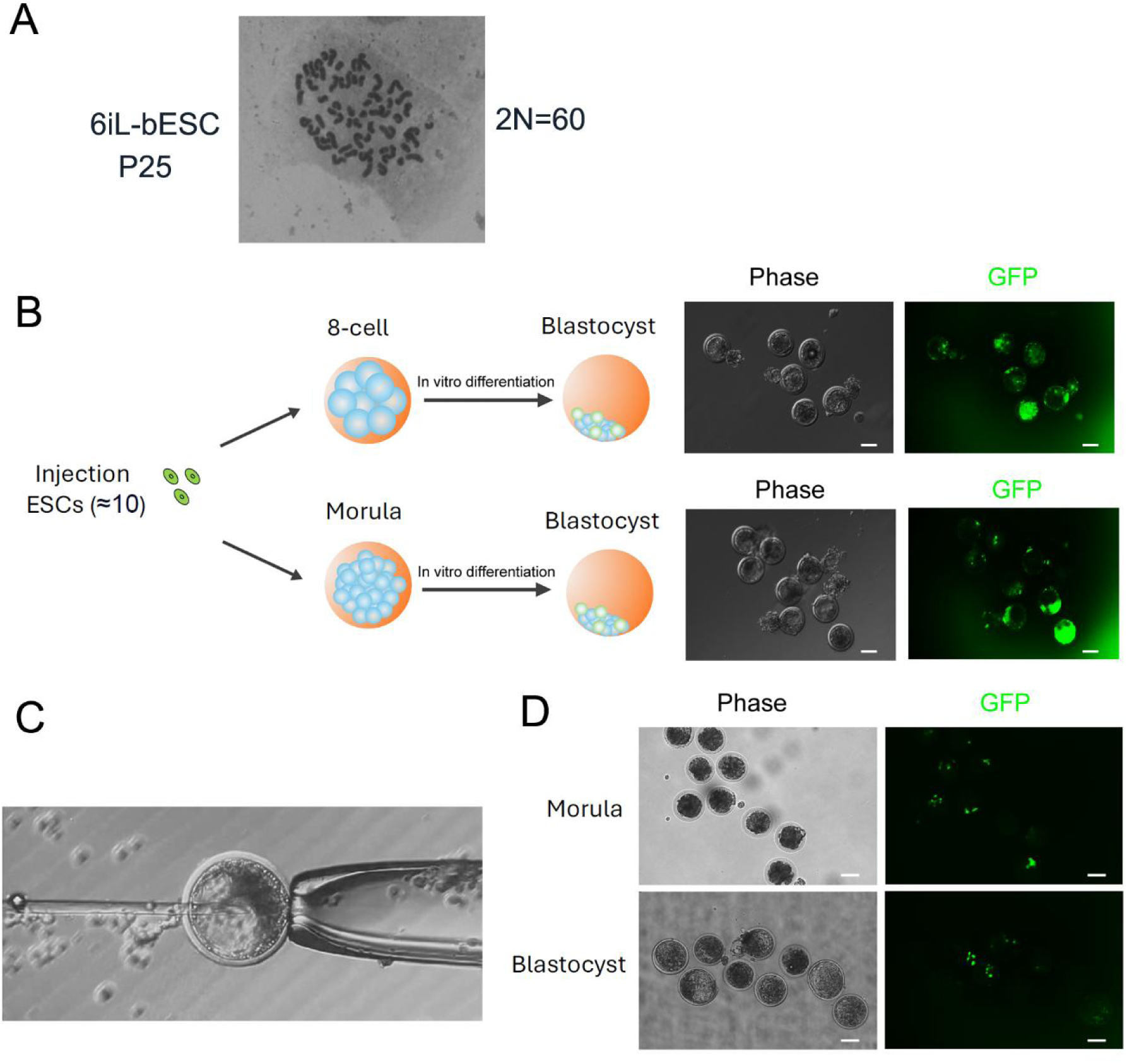
Chimeric Contribution of bESCs in Bovine Embryos. (A) Karyotyping of bESC at passage 25. (B) Schematic representation of the injection of GFP-labeled ESCs (8-10 cells) into 8-cell and morula-stage bovine embryos, followed by in vitro culture to the blastocyst stage. Representative phase-contrast and GFP FL images show the presence of GFP-positive cells in blastocysts. Scale bars, 100 µm. (C) Microinjection of GFP-labeled ESCs into a bovine blastocyst. (D) Representative phase-contrast and GFP fluorescence images showing GFP-positive ESCs in morula- and blastocyst-stage embryos following injection and in vitro culture. Scale bars, 100 µm.

**Figure S5.**
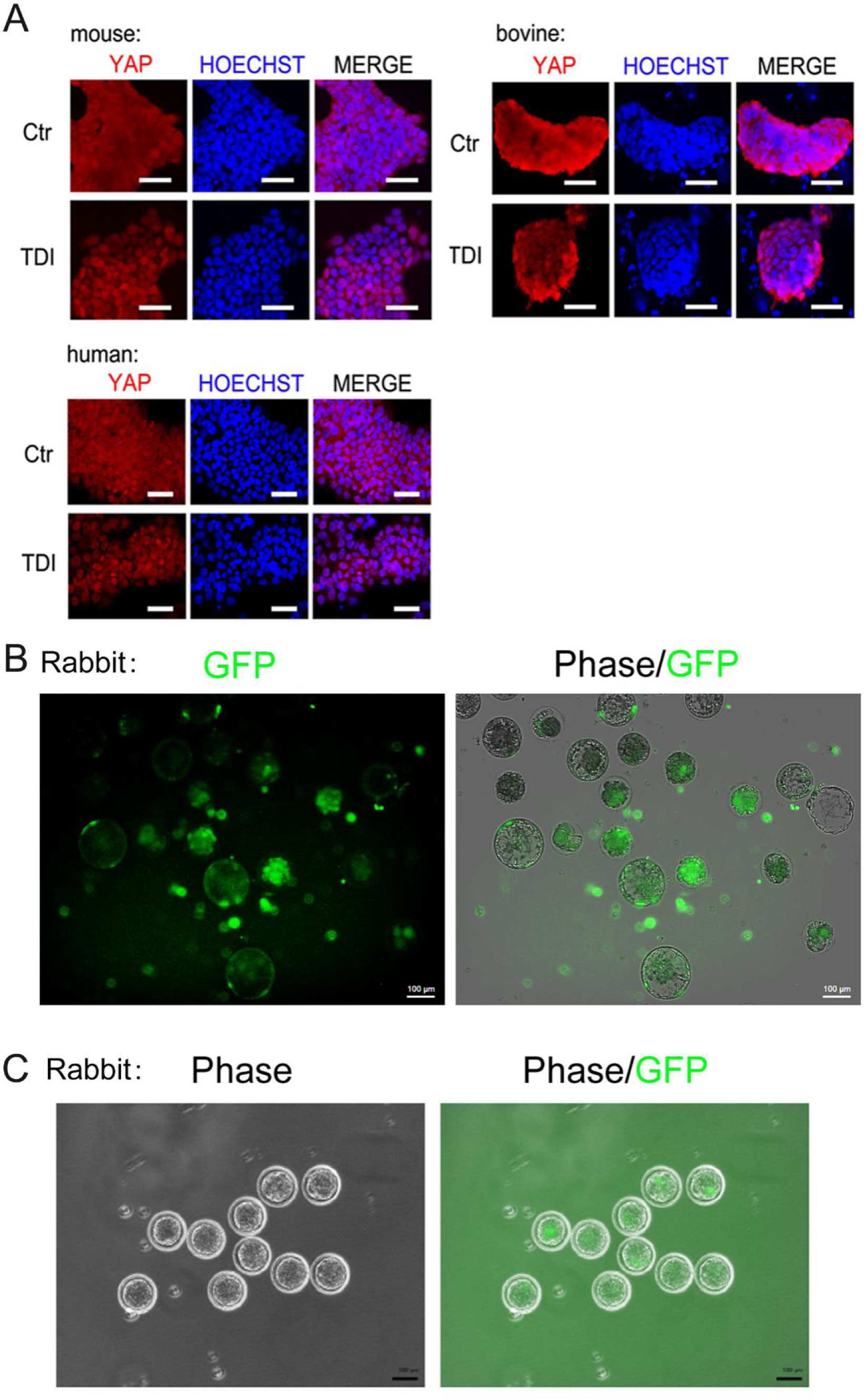
Distribution of GFP-labeled 6iL-rabESCs in morula and blastocyst-stage embryos. (A) IF staining of YAP (red) and nuclear marker HOECHST (blue) in mouse, bovine, and human ESCs under control (Ctr) and TDI conditions. Scale bars: 50 µm. (B)(C) GFP-labeled 6iL-rabESCs were injected into 8-cell stage rabbit embryos and allowed to develop to the morula and blastocyst stages in vitro. FL results indicate the localization of GFP-injected cells. (B) Blastocyst or morula-stage rabbit embryos showing GFP-positive cells. Scale bar: 100 μm. (C) Morula-stage embryos with GFP-positive cells. Scale bar: 100 μm.

**Figure S6.**
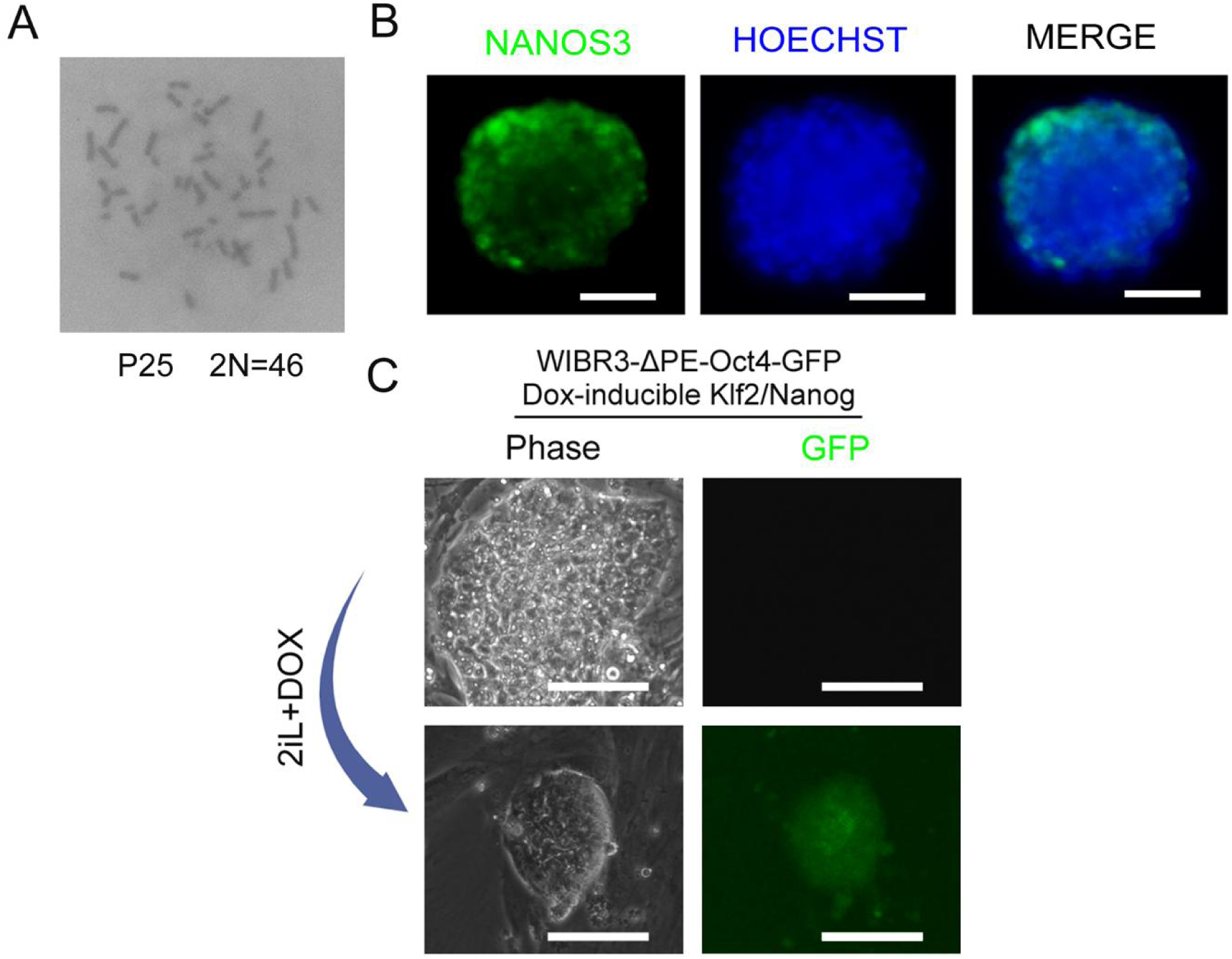
Characteristics of Human Naive ESCs Cultured Under 6iL Conditions. (A) Karyotype analysis of passage 25 (P25) 6iL human iPSCs, showing a normal diploid chromosome number (2N=46). (B) Representative IF analysis of analysis of PGC-LCs differentiated from 6iL-hiPSCs, showing expression of PGC-specific proteins NANOS3 (green) and HOECHST (blue). Scale bars = 50 µm. (C) WIBR3-ΔPE-Oct4-GFP cells carrying Dox-inducible Klf2/Nanog were cultured under 2iL +DOX conditions. Phase contrast (left) and GFP fluorescence (right) images are shown. Scale bars=100µm.

**Table S1.** List of qPCR primers.

**Table S2.** List of 6iL ESC derivation efficiency.

**Movie S1.** Beating cardiomyocytes differentiated from 6iL-rabESCs.

